# Developmental regulators drive DUX4 expression in facioscapulohumeral muscular dystrophy

**DOI:** 10.1101/2024.05.21.595131

**Authors:** Amelia Fox, Jonathan Oliva, Rajanikanth Vangipurapu, Francis M. Sverdrup

**Author notes:** Corresponding Author: Francis M. Sverdrup;.

## Abstract

Facioscapulohumeral muscular dystrophy (FSHD) is a progressive muscle wasting disease caused by misexpression of the Double Homeobox 4 (DUX4) transcription factor in skeletal muscle. While epigenetic derepression of D4Z4 macrosatellite repeats is recognized to cause DUX4 misexpression in FSHD, the factors promoting *DUX4* transcription are unknown. Here, we show that SIX (*sine oculis*) transcription factors, critical during embryonic development, muscle differentiation, regeneration and homeostasis, are key regulators of *DUX4* expression in FSHD muscle cells. In this study, we demonstrate SIX1, SIX2, and SIX4 to be necessary for induction of *DUX4* transcription in differentiating FSHD myotubes *in vitro*, with SIX1 and SIX2 being the most critical in driving *DUX4* expression. Interestingly, DUX4 downregulates *SIX* RNA levels, suggesting negative feedback regulation. Our findings highlight the involvement of SIX transcription factors in driving the pathogenesis of FSHD by promoting *DUX4* and DUX4 target gene expression.

**Teaser:** We identified a family of developmental regulators that promote aberrant *DUX4* expression in FSHD differentiating muscle cells.

## Introduction

Double Homeobox 4 (DUX4) is a transcription factor with a critical role during early embryonic development and when ectopically expressed in the pathogenesis of several diseases (Mocciaro et al. 2021; Campbell et al. 2018). During normal development, DUX4 is expressed in two- and four-cell stage embryos and activates the zygotic genome transcriptional program by inducing the transcription of hundreds of genes (Mocciaro et al. 2021; De Iaco et al. 2017). Subsequently, *DUX4* expression is downregulated at the 8-cell stage and silenced in most somatic tissues by repeat-mediated epigenetic repression (van Overveld et al. 2003; Snider et al. 2010). However, loss of repression and *DUX4* activation by unknown transcriptional and epigenetic regulators drives ectopic *DUX4* expression, leading to one of the most common muscular dystrophies, facioscapulohumeral muscular dystrophy (FSHD) (Mocciaro et al. 2021; Smith et al. 2023; Deenen et al. 2014). FSHD, the third most prevalent muscular dystrophy, is characterized by progressive skeletal muscle weakness and asymmetric muscle wasting, often first noticed in the face, shoulders, and upper arms (Campbell et al. 2018; Tawil and Van Der Maarel 2006). Bursts of *DUX4* expression in the diseased state leads to progressive muscle degeneration, inflammation, fat infiltration and inadequate muscle regeneration (Mocciaro et al. 2021; Banerji et al. 2020; Kan et al. 2010). Importantly, aberrant DUX4 expression has also been implicated in several cancers, where its expression leads to MHC class I antigen suppression, immune evasion and loss of checkpoint blockades, resulting in cancer progression and immunotherapy failure (Smith et al. 2023; Mocciaro et al. 2021; Jongsma, Neefjes, and Spaapen 2021; Chew et al. 2019). Therefore, defining the factors regulating *DUX4* transcriptional activation is critical in understanding DUX4- associated disease development and progression.

There are two main classifications of FSHD (FSHD1 and FSHD2), with clinically identical symptoms, that are distinguished by genetic differences (Statland and Tawil 2014; Lemmers et al. 2010). FSHD1, affecting most patients, results from a pathogenic contraction of the D4Z4 macrosatellite repeat array (1- 10 repeats compared to 11-100 repeats in unaffected persons) at the subtelomeric region of chromosome 4q35 (Tawil, van der Maarel, and Tapscott 2014; Himeda and Jones 2019; van der Maarel et al. 2012; Wijmenga et al. 1990). FSHD2, however, develops from loss-of-function mutations in one or more chromatin modifiers (e.g., SMCHD1, DNMT3B, LRIF1) leading to decreased DNA methylation and chromatin relaxation at D4Z4 repeats (Himeda and Jones 2019; Hamanaka et al. 2020; van der Maarel et al. 2012; Balog et al. 2015). Consistent between FSHD1 and FSHD2, disease penetrance, onset, progression and severity vary considerably between patients (Tawil, van der Maarel, and Tapscott 2014; van der Maarel, Frants, and Padberg 2007; Jones et al. 2012). Most patients are diagnosed within their second decade of life (Himeda and Jones 2019). Patients exhibiting clinical symptoms before the age of 10 are classified as having early disease-onset and exhibit a more rapidly progressing and severe disease state (Brouwer et al. 1994; Goselink et al. 2018; Statland and Tawil 2014). Developmental comorbidities associated with early disease-onset include hearing loss and retinal vasculopathy and are linked to having a greater contraction of D4Z4 (1-3 residual repeats) (Padberg et al. 1995; Lutz et al. 2013; Brouwer et al. 1991). Due to DUX4 being the common link between FSHD1 and FSHD2, targeting *DUX4* mRNA has been recognized as an attractive therapeutic target (Himeda, Jones, and Jones 2015; Oliva et al. 2019). However, the factors responsible for driving *DUX4* expression in skeletal muscle remain unknown.

SIX (*sine oculis*) transcription factors are part of an established and evolutionarily conserved network of transcription factors known as the PSED (<u>P</u>AX-<u>S</u>IX-<u>E</u>YA-<u>D</u>ACH) network (Maire et al. 2020; Viaut, Weldon, and Munsterberg 2021; Kumar 2009). Together, this network has critical regulatory roles in the development and regeneration of many tissues including most sensory organs, the kidney, and skeletal muscle (Viaut, Weldon, and Munsterberg 2021; Maire et al. 2020; Inoue et al. 2023; Kumar 2009; Yajima et al. 2010). There are three distinct subclasses of SIX transcription factors that are grouped by similarities in their amino acid sequences: SIX1/2 (*sine oculis*), SIX3/6 (*optix*), and SIX4/5 (*DSIX4*) (Kumar 2009; Seo et al. 1999; Maire et al. 2020). However, only SIX1, SIX2, SIX4, and SIX5 are expressed in embryonic myogenesis, proliferating myogenic stem cells and adult myofibers (Maire et al. 2020; Wurmser et al. 2023). During muscle development, these transcription factors function by controlling myogenic regulatory factors (MRFs) to direct myogenic cell fate decisions, muscle differentiation and muscle regeneration through stem cell renewal (Maire et al. 2020; Le Grand et al. 2012). During these processes, SIX transcription factors can function as transcriptional activators or repressors independently and/or in association with a cofactor, such as EYA (transcriptional activator) or DACH (transcriptional repressor) (Maire et al. 2020). Importantly, when absent or ectopically expressed, SIX transcription factors and their related cofactors are responsible for several congenital disorders (e.g., BOR syndrome, hearing loss and craniofacial abnormalities) and cancers by promoting increased invasion and metastasis (Meurer et al. 2021; Ruf et al. 2004; Wu et al. 2015). Although the roles of SIX transcription factors in development and cancer have been extensively studied, their potential contributions in muscular dystrophies, like FSHD, have yet to be defined. In this study, we elucidate the function of SIX transcription factors in the regulation of *DUX4* transcription in FSHD.

## Results

### Knockdown of SIX1, SIX2 and SIX4 suppresses DUX4 expression in differentiating FSHD muscle cells

DUX4 is a pioneer transcription factor that can bind heterochromatic genomic regions to transcriptionally activate hundreds of downstream targets, leading to the FSHD phenotype (Himeda and Jones 2019; van der Maarel, Frants, and Padberg 2007; van der Maarel, Tawil, and Tapscott 2011).

*DUX4* is induced during early myogenic differentiation of FSHD myoblasts, *in vitro* (Bosnakovski et al. 2018; Balog et al. 2015); however, the transcriptional components that activate *DUX4* expression in skeletal muscle and other non-muscle tissues associated with FSHD are undefined. To identify transcription factors that drive bursts of *DUX4* expression in FSHD myotubes, we began screening factors critical during muscle development and myogenic differentiation using small interfering RNA (siRNA)-mediated knockdown in immortalized patient-derived FSHD1 (54-2 and 16-ABIC) and FSHD2 (MB200) myoblast lines. We focused on the SIX transcription factor family after identifying novel phosphorylation sites on SIX4 that appear to be sensitive to p38 inhibition. This phosphorylation profile is of interest due to p38 MAPK involvement in *DUX4* regulation (Oliva et al. 2019). We transfected myoblasts with siRNAs selectively targeting SIX genes (SIX1, SIX2, and SIX4) prior to inducing differentiation to generate multinucleated myotubes. Quantitative real-time PCR (qPCR) was used to quantify *DUX4* and DUX4 target mRNA levels. We found that selective knockdown of SIX1, SIX2 or SIX4, individually, led to significant decreases in *DUX4* mRNA levels in FSHD1 (54-2) and FSHD2 (MB200) myotubes, with *SIX1* and *SIX2* knockdown having the most pronounced effects on *DUX4* levels (Fig. 1A and fig. S1A). DUX4 targets (*MBD3L2*, *LEUTX*, and *ZSCAN4)* were also significantly reduced, largely mirroring the decreases in *DUX4* mRNA (Fig. 1A, fig. S1A and fig. S2A). Based on the relative expression levels from qPCR, we determined *SIX5* mRNA was nearly undetectable in our cell system (table S1). These results suggest the individual involvement of primarily SIX1, SIX2 and SIX4 in the transcriptional activation of *DUX4* gene expression in FSHD myotubes.

**Figure 1.**
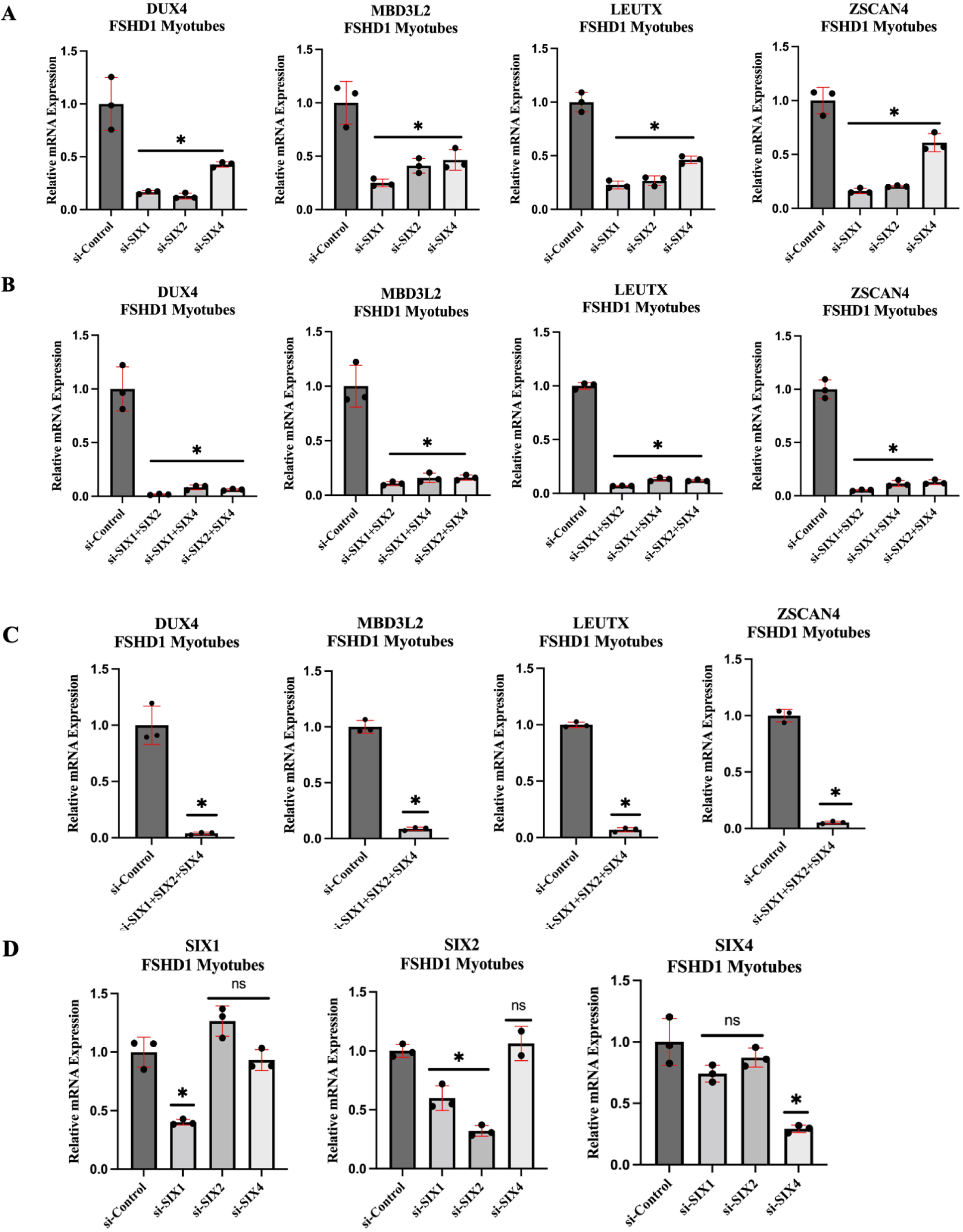
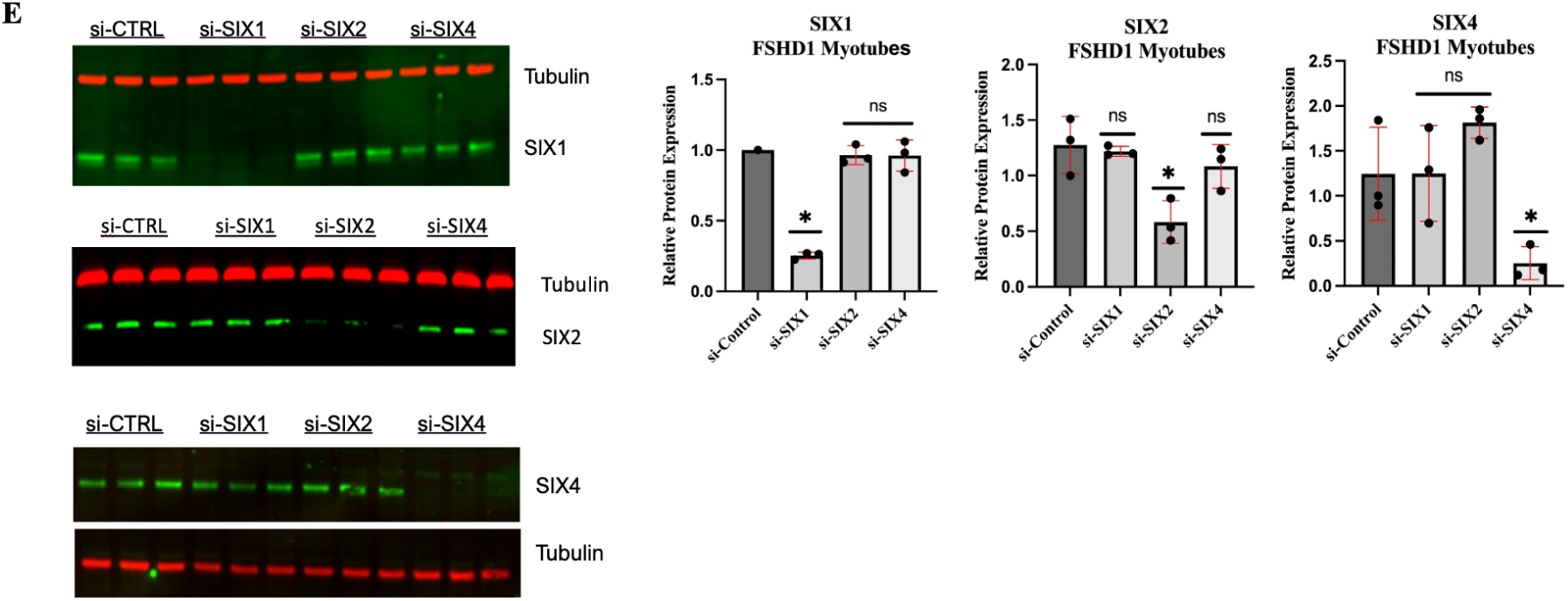
siRNA knockdown of SIX1, SIX2 and SIX4 suppresses DUX4 mRNA in FSHD1 patient-derived myotubes. FSHD1 (54-2) myoblasts were transfected with siRNA targeting SIX1, SIX2, and SIX4 individually (**A**) and combinatorially (**B-C**) and differentiated to form multinucleated myotubes. Shown are the relative mRNA levels for *DUX4* and DUX4 targets (*MBD3L2*, *LEUTX,* and *ZSCAN4*) (**A-C**) and each *SIX* paralog (**D**). (**E**) Western blots show individual protein depletion of SIX1, SIX2 and SIX4. Alpha-tubulin is shown as the loading control. Western blots were quantified to demonstrate relative knockdown of each protein. Each experimental condition (n=3) was normalized to a negative si-Control with the mean and standard deviation depicted. Asterisks demonstrate statistical significance between the control vs experimental group using an unpaired two tailed *t-*test (ns= not significance; *p<0.05).

Next, we knocked down various combinations of *SIX*, *SIX2*, and *SIX4* to identify potential non-redundant roles between each paralog and to determine if we could further suppress *DUX4* mRNA levels. Dual combinations containing *SIX2* siRNA had the most significant reduction of *DUX4* and DUX4 target transcripts with the combined knockdown of *SIX1* and *SIX2* eliciting the greatest decrease in *DUX4* and target gene mRNA levels (∼98% decrease of *DUX4* mRNA) (Fig. 1B, fig. S1B, and fig. S2B). Combined knockdown of all three SIX genes resulted in ∼90-98% decrease in *DUX4* and DUX4 target gene expression in FSHD1 (54-2 and 16-ABICs) and FSHD2 (MB200) myotubes (Fig. 1C, fig. S1C, and fig. S2C). siRNAs selectively targeting *SIX1*, *SIX2*, and *SIX4* effectively knockdown their respective target mRNA levels in both FSHD1 and FSHD2 myotubes (Fig. 1D and fig. S1D). Western blotting analysis and quantification confirmed selective protein depletion of SIX1, SIX2 and SIX4 by their respective siRNAs (Fig. 1E). Together, these data suggest that expression of SIX transcription factors (SIX1, SIX2, and SIX4) are required and combinatorially regulate *DUX4* transcription, with SIX1 and SIX2 acting as the main drivers of DUX4 expression.

### Combined knockdown of SIX1, SIX2, and SIX4 does not inhibit myogenic differentiation

SIX proteins are well documented to participate in muscle differentiation and regeneration through their control of myogenic regulatory factors (MRFs) (Maire et al. 2020). To rule out the possibility that knockdown of SIX proteins indirectly affected *DUX4* expression by inhibiting myogenic differentiation, we monitored myotube morphology and measured myogenic markers after the combined knockdown of SIX1, SIX2, and SIX4 in differentiating FSHD1 (54-2 and 16-ABIC) and FSHD2 (MB200) cells. Consistent between the tested cell lines, we observed no delays in myotube formation, morphology or fusion index of the generated multinucleated myotubes in comparison to cells treated with negative control siRNA (Fig. 2A-C, fig. S1E-G, and fig. S2D). We also measured gene expression of several myogenic markers. We screened markers of myogenic commitment (myoblasts determination protein 1, *MYOD1*), early differentiation (myogenin, *MYOG*), and late differentiation (creatine kinase M-Type, *CKM*). Additionally, myosin heavy chains involved in embryonic development and regeneration (myosin heavy chain, *MYH3* and *MYH8*), slow fiber types (myosin heavy chain 7, *MYH7*), and fast fiber types (myosin heavy chains, *MYH1*, *MYH2*, and *MYH4*) were analyzed. qPCR analysis revealed that the combined knockdown of *SIX1*, *SIX2*, and *SIX4* had no differential effects on *MYOD*, *MYOG*, *MYH1* and *MYH4* mRNA levels, suggesting that knockdown of SIX proteins does not generally inhibit early myotube differentiation (Fig. 2D and Fig. 2E). However, there were differential effects on *CKM* (decreased), *MYH2* (increased), *MYH7* (decreased), *MYH3* (decreased), and *MYH8* (decreased) with the combined knockdown of all three SIX genes (Fig. 2D-G). These results suggest that the effects of SIX1/2/4 knockdown on *DUX4* expression, *in vitro,* are not due to inhibition of differentiation, but instead related to specific regulation of target genes.

**Figure 2.**
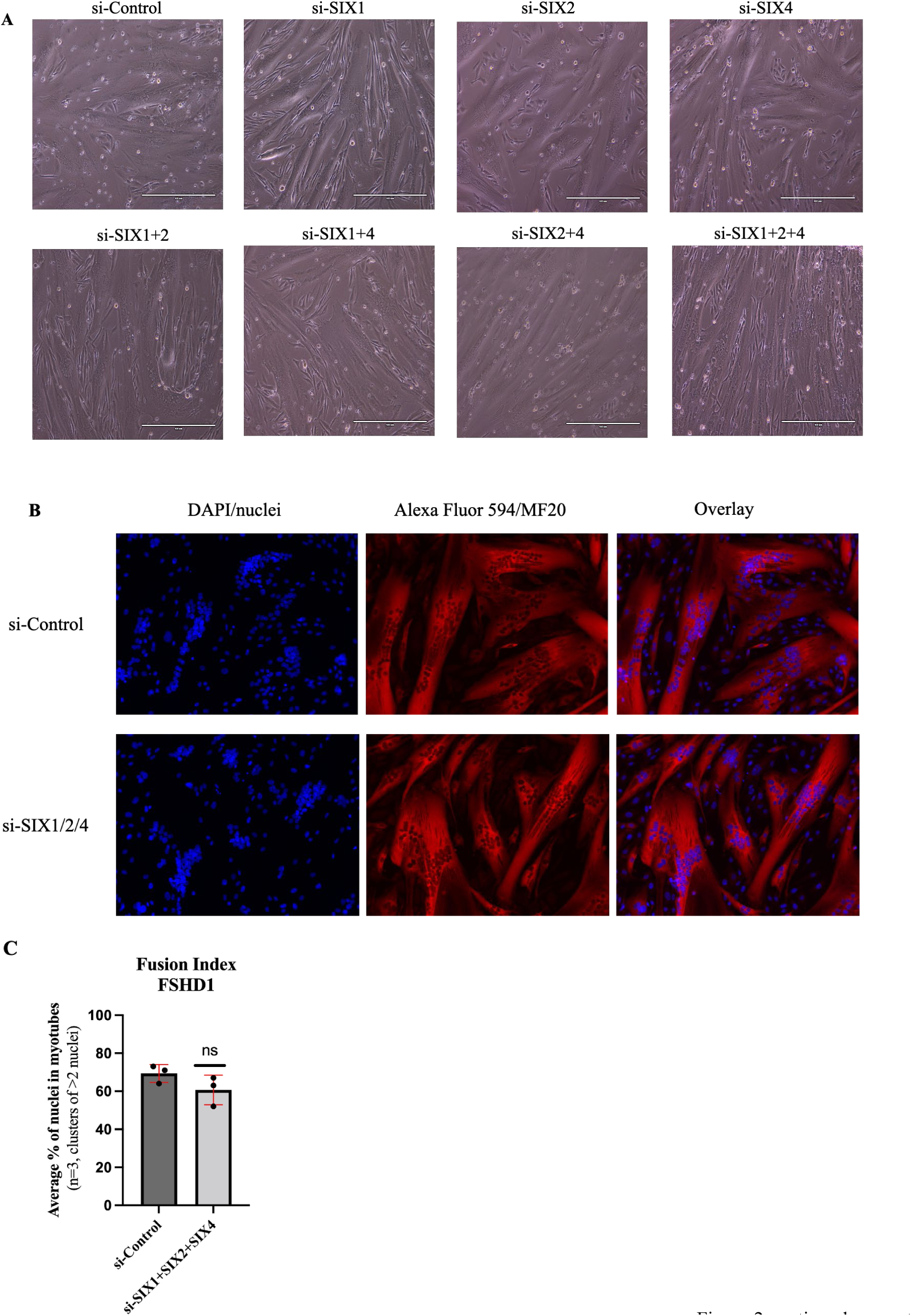
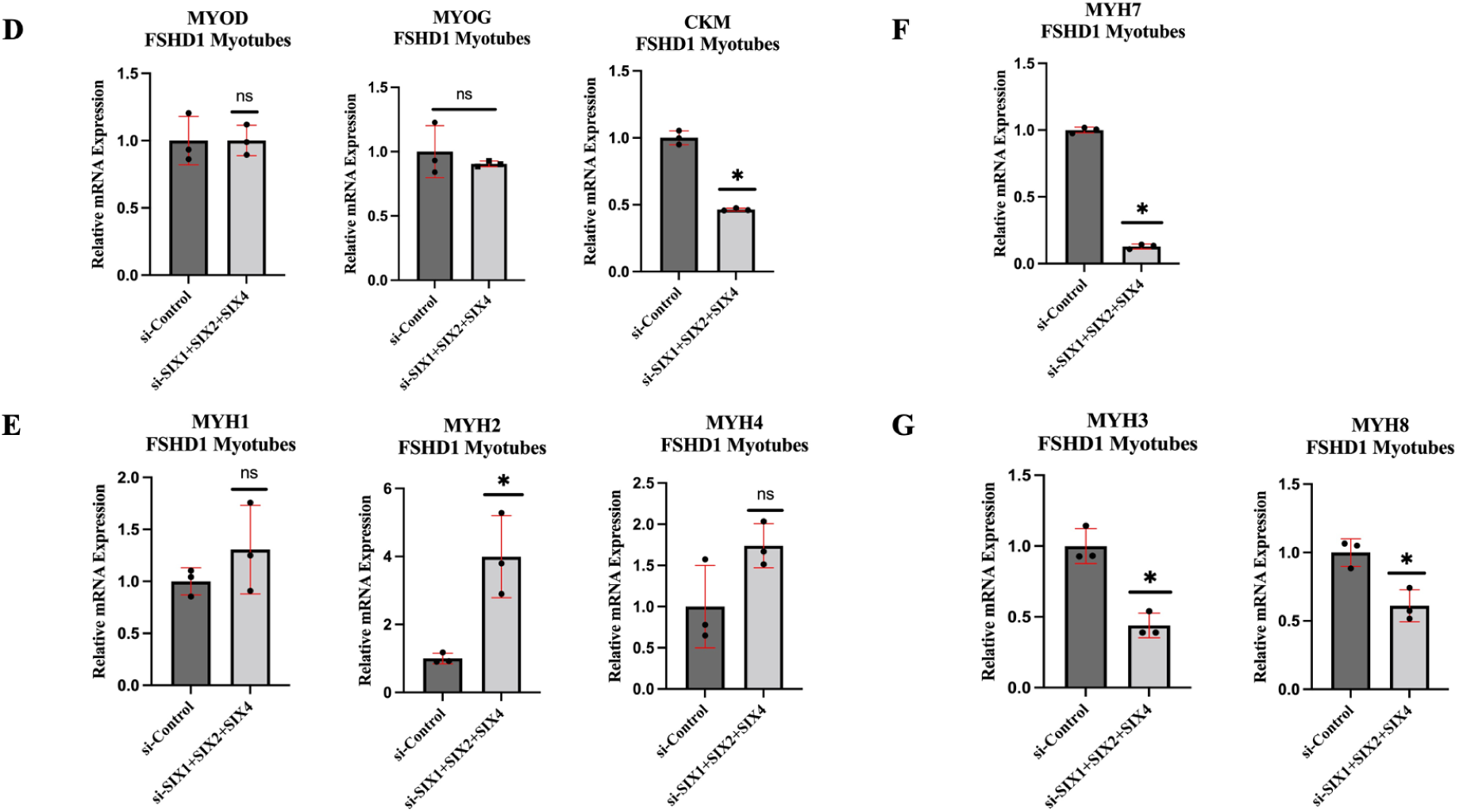
Combined knockdown of SIX1, SIX2, and SIX4 does not inhibit myogenic differentiation. FSHD1 (54-2) myoblasts were transfected with siRNAs targeting SIX1, SIX2 and SIX4, and differentiated to generate multinucleated myotubes. (**A**) Brightfield images showing the morphology of multinucleated myotubes with clusters of nuclei for the individual and combined knockdown(s) of SIX1/2/4. (**B**) Immunofluorescence staining for myosin heavy chain (MF20, red) with counterstaining of nuclei (blue) for si-Control and si-SIX1/2/4 conditions. (**C**) Fusion index was calculated (n=3) for the conditions shown in **B**. (**D-G**) mRNA levels for markers for myogenic commitment (MYOD), early (MYOG) and late (CKM) differentiation (**D**), fast fiber-types (**E**), slow fiber-types (**F**) and regeneration (**G**). Each experimental condition (n=3) was normalized to a negative si-Control with the mean and standard deviation depicted. Asterisks demonstrate statistical significance between the control vs experimental group using an unpaired two tailed *t-*test (ns= not significance; *p<0.05).

### EYA coactivators participate in activating *DUX4* transcription

Since SIX1 (as opposed to SIX2 and SIX4) does not possess an intrinsic transcriptional activation domain and requires coactivators to positively affect transcription (Blevins et al. 2015), we knocked down the most highly expressed EYA genes (*EYA1*, *EYA3*, and *EYA4*) in our FSHD myoblast lines to assess their potential involvement in *DUX4* regulation. We found that the combined knockdown of EYA1, EYA3 and EYA4 resulted in a 54% decrease of *DUX4* and a 40-70% decrease of DUX4 target mRNA levels upon myogenic differentiation of FSHD1 (54-2) myoblasts (Fig. 3A and Fig. 3C).

**Figure 3.**
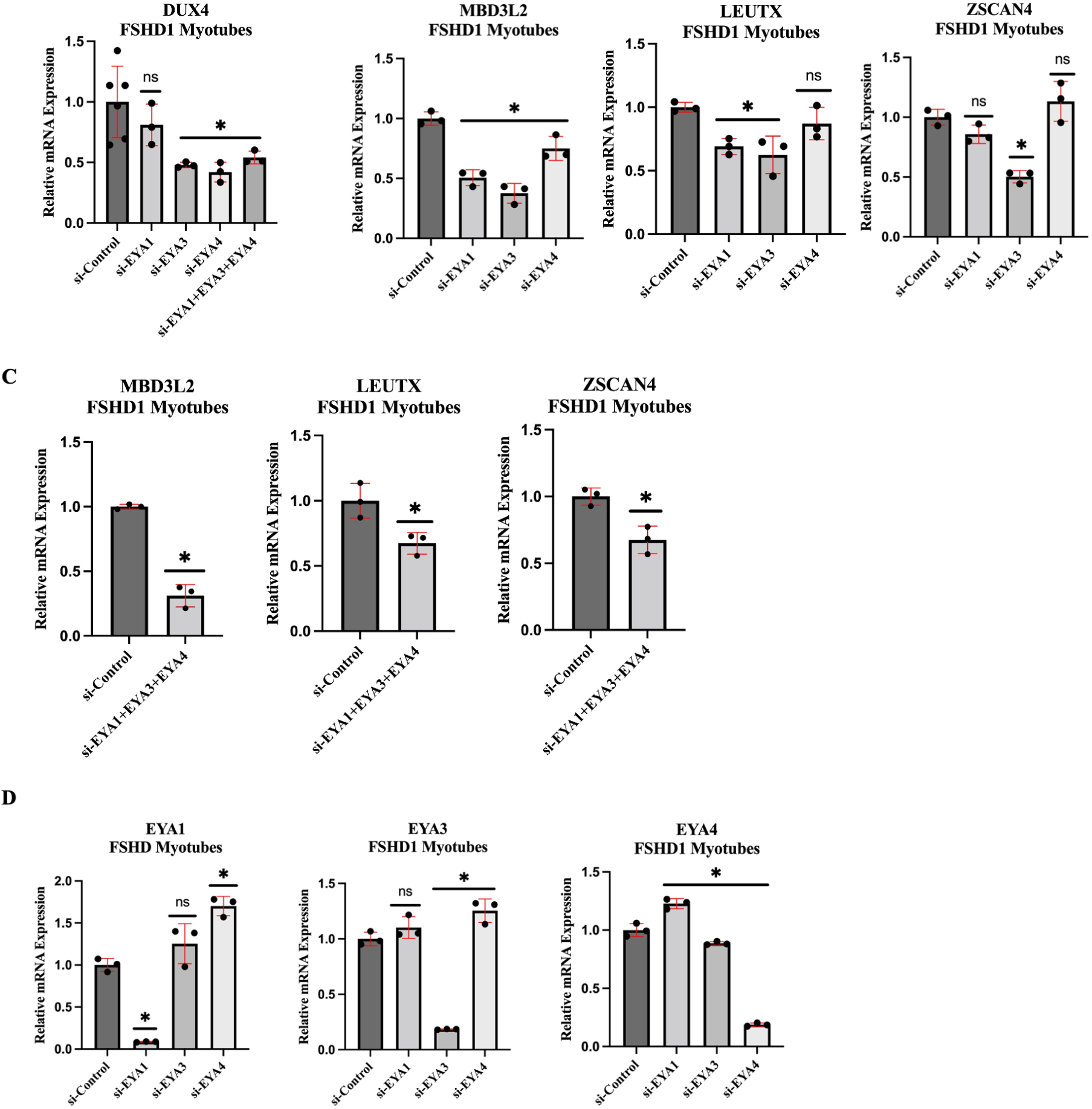
Involvement of SIX coactivator, EYA, in DUX4 regulation. FSHD1 (54-2) myoblasts were transfected with EYA1, EYA3 and EYA4 and differentiated to form multinucleated myotubes. Shown are the relative mRNA levels from qPCR for *DUX4* (**A**), DUX4 targets (*MBD3L2*, *LEUTX* and *ZSCAN4*) (**B** and **C**) and the relative knockdown for each EYA target (**D**). Each experimental condition (n=3) was normalized to a negative si-Control with the mean and standard deviation depicted. Asterisks demonstrate statistical significance between the control vs experimental group using an unpaired two tailed *t-*test (ns= not significance; *p<0.05).

Individual knockdown of EYA1 did not significantly affect *DUX4*, but knockdown of EYA3 or EYA4 resulted in significant *DUX4* decreases with EYA3 having the most pronounced effect (Fig. 3A-C and fig. S3A). siRNAs selectively targeting *EYA1*, *EYA3* and *EYA4* successfully knocked down each of their intended targets (Fig. 3D). This data is consistent with EYA coactivators playing some role in promoting DUX4 expression through their interaction with SIX transcription factors but leaves open the possibility that mechanisms independent of EYA coactivation are important.

### Knockdown of SIX proteins suppresses DUX4 expression in a differentiation-dependent manner

Having established the requirement of SIX transcription factors to drive differentiation-mediated bursts of *DUX4* expression, we tested whether siRNA knockdown of SIX transcription factors would also suppress the relatively low levels of DUX4 expression in undifferentiated myoblasts (Bosnakovski et al. 2018; Cowley et al. 2023; Tassin et al. 2013; Rickard, Petek, and Miller 2015). We transfected FSHD1 (54-2) and FSHD2 (MB200) myoblasts with *SIX* siRNAs and maintained the cells for 72 hours prior to qPCR analysis to allow for sufficient protein turnover. Individual and combined knockdown of *SIX1*, *SIX2*, and *SIX4* showed no significant effects on the low level of *DUX4* and target mRNA levels in FSHD1 and FSHD2 myoblasts (Fig. 4A-D and fig. S4A). To determine if SIX transcription factors were otherwise active in myoblasts, we looked at the expression of known targets of SIX1 (*PGK1, SLC4A7*) and SIX2 (*EYA1*, *SLC4A7*) (Gu et al. 2016; Li et al. 2018; Jin et al. 2021). Knockdown of *SIX1* resulted in a 26% decrease in *PGK1* RNA levels and a 46% decrease in *SLC4A7* RNA levels (Fig. 4E), indicating that SIX1 does exhibit transcriptional activity towards known targets in myoblasts.

**Figure 4.**
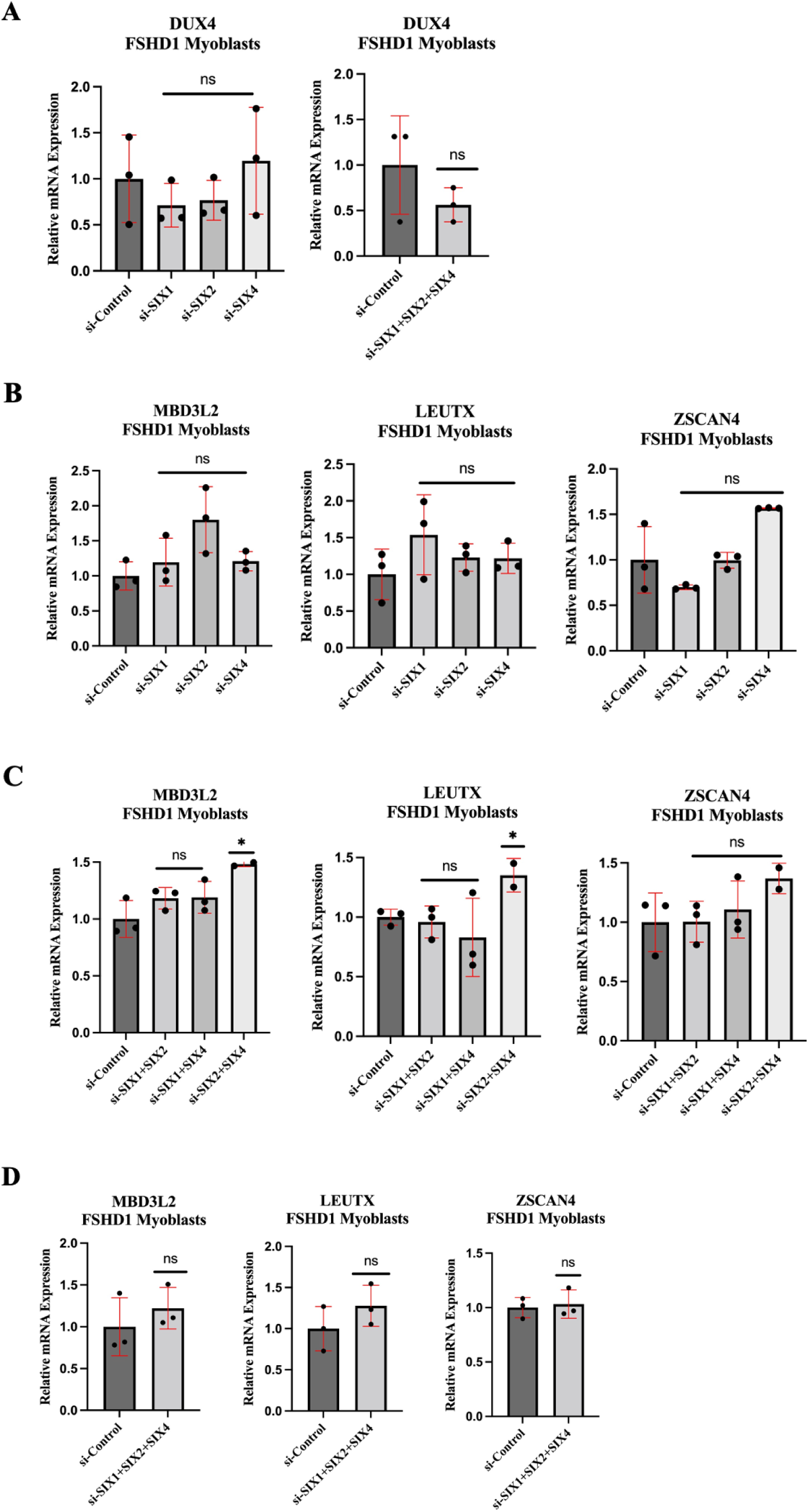
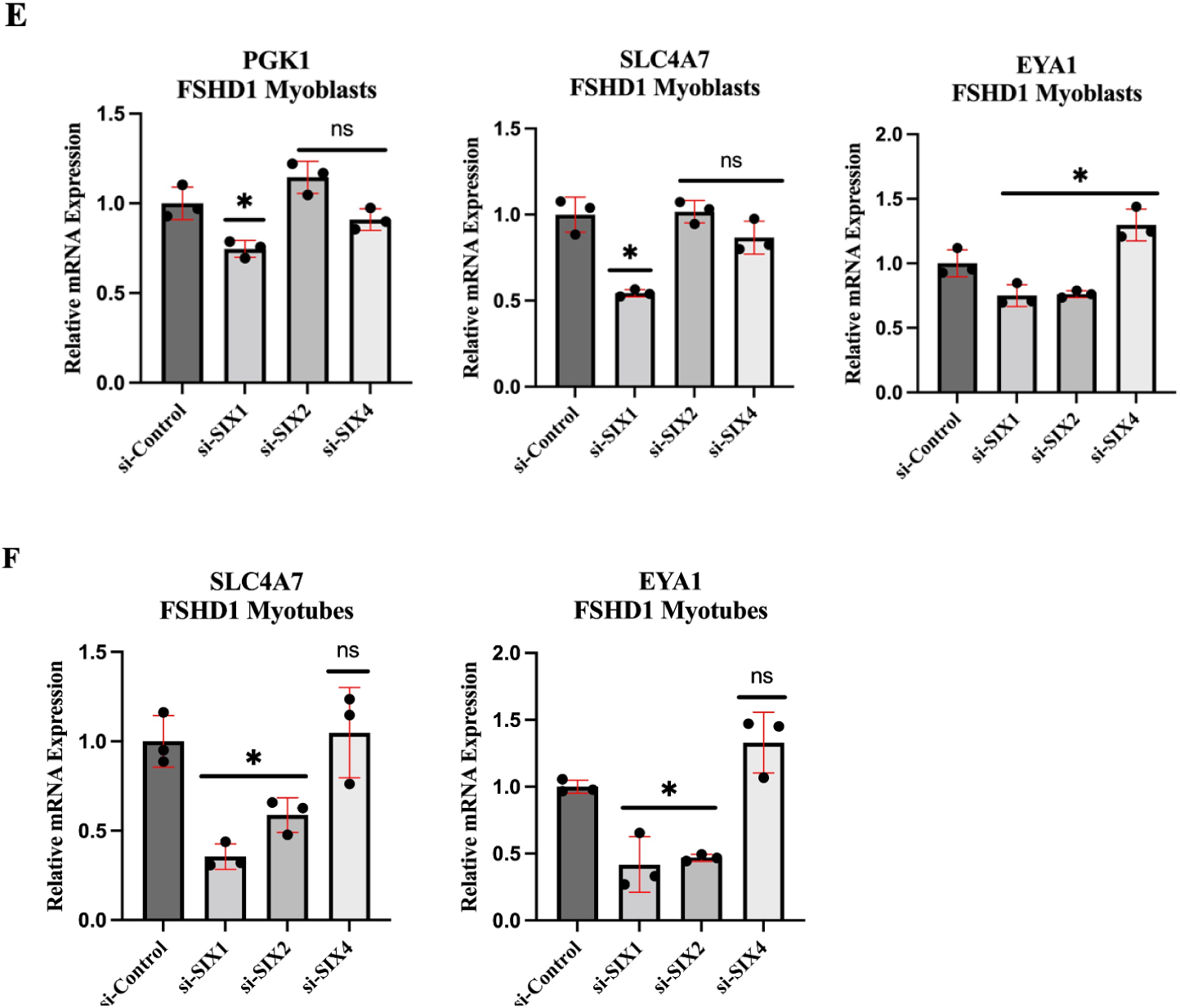
Low level of DUX4 expression in myoblasts are unaffected by SIX1/2/4 knockdown. FSHD1 (54-2) were transfected with SIX1, SIX2 and SIX4 siRNAs in proliferating myoblasts. Relative mRNA levels for *DUX4* (**A**) and DUX4 targets (**B-D**) for individual and combined SIX1/2/4 knockdown. Effects of SIX1 and SIX2 knockdown on mRNA levels for targets (*PGK1*, *SLC4A7* and *EYA1*) in myoblasts (**E**) and myotubes (**F**). Each experimental condition (n=3) was normalized to a negative si-Control with the mean and standard deviation depicted. Asterisks demonstrate statistical significance between the control vs experimental group using an unpaired two tailed *t-*test (ns= not significance; *p<0.05).

Conversely, knockdown of *SIX2* in myoblasts resulted in a modest 25% decrease in *EYA1* RNA levels but failed to decrease *SLC4A7* RNA (Fig. 4E). In differentiating myotubes, however, knockdown of *SIX1* and *SIX2* resulted in a more substantial decrease in EYA1 levels (54-59% decrease) and significantly decreased *SLC4A7* RNA levels (42-64% decrease) (Fig. 4F). Taken together, these data indicate that the regulation of *DUX4* expression by SIX1, SIX2, and SIX4 occurs primarily during the differentiation-induced increases of *DUX4*, and that while SIX1 and SIX2 have demonstrable transcriptional activity in myoblasts, their ability to regulate DUX4 requires myogenic differentiation signaling.

### SIX1 and SIX2 do not govern restricted DUX4 expression to a subset of nuclei

*DUX4* transcription is restricted to a rare subset of nuclei in differentiating FSHD myotubes, resulting in clusters of adjacent nuclei importing DUX4 protein due to the syncytial nature of multinucleated myotubes, yet the reason for this restriction is not understood (Talbot and Maves 2016; Yao et al. 2014; van der Maarel, Tawil, and Tapscott 2011). One possibility is the restricted expression of necessary transcription factors that drive *DUX4* transcription. To determine if the restricted transcription of *DUX4* in a subset of nuclei was due to restricted expression of SIX proteins in the same nuclei, we stained for SIX1 and SIX2, due to their prominent role in regulating *DUX4* transcription. Immunofluorescence staining in FSHD1 cells revealed SIX1 and SIX2 protein to be present in every nucleus in proliferating myoblasts and along multinucleated myotubes (Fig. 5A-D). Staining intensity was significantly decreased with siRNA knockdown of SIX1 and SIX2, and increased with overexpression of SIX1 and SIX2, confirming specificity of staining (Fig. 5A-D). These results indicate that SIX1 and SIX2 expression is ubiquitous and does not explain the restricted expression of DUX4 to a subset of muscle cell nuclei in FSHD.

**Figure 5.**
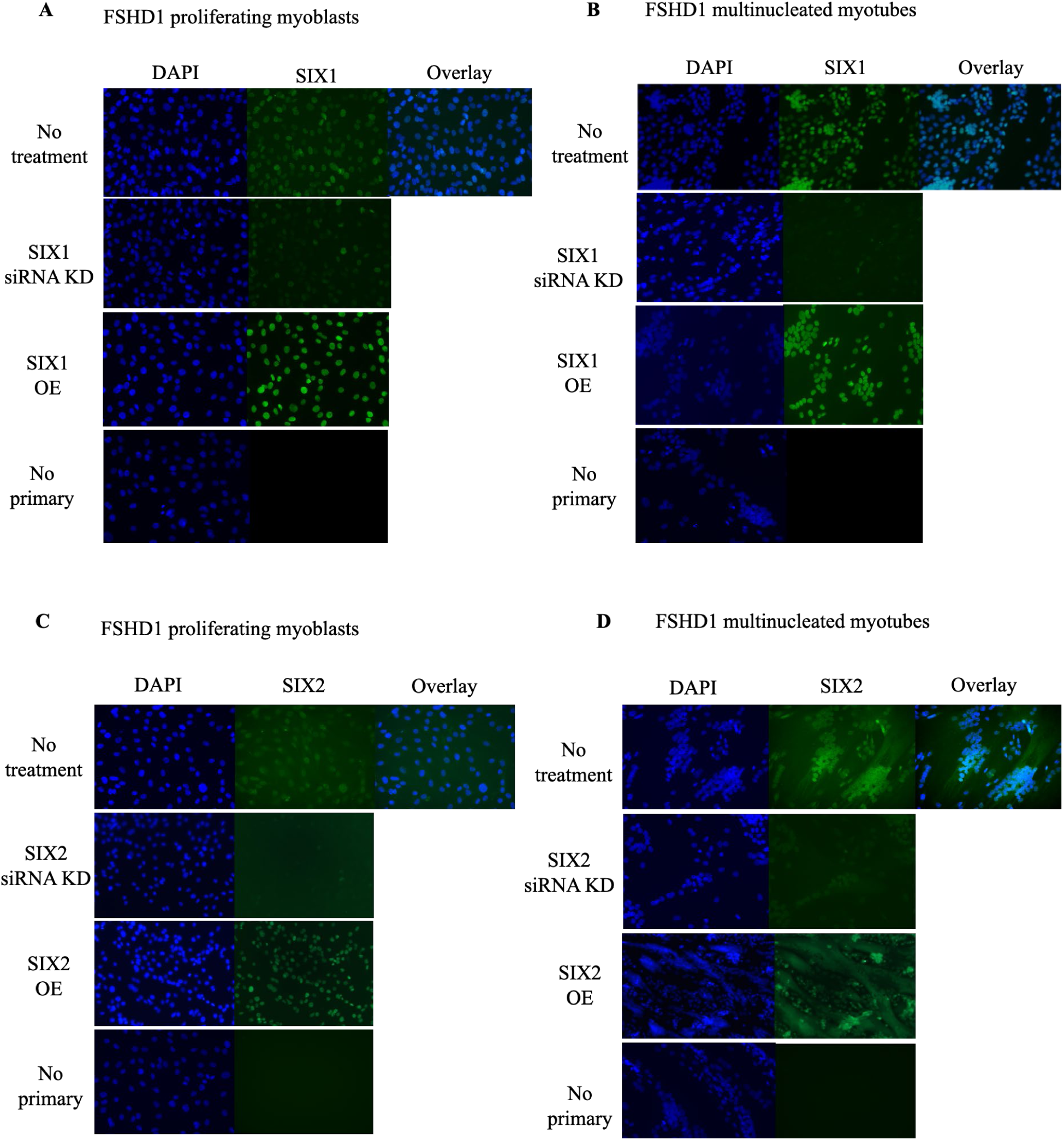
SIX1 and SIX2 protein expression in FSHD myoblasts and myotubes. FSHD1 (54-2) proliferating myoblasts (left) and myotubes (right) were stained with anti-SIX1 (**A** and **B**) and anti- SIX2 (**C** and **D**) antibodies (green) and nuclei counterstained with DAPI (blue).

### SIX1 and SIX2 levels are not limiting for DUX4 transcription

We tested whether increased expression of SIX1 or SIX2 would further induce DUX4 during differentiation. Stable FSHD myoblasts lines overexpressing SIX1 or SIX2 were created using lentivirus vectors. Western blot analysis confirmed a 13-fold increase in myoblasts and 7-fold increase in myotubes in steady state SIX1 protein levels and a 10-fold increase in SIX2 protein levels in transduced FSHD1 (54-2) myoblasts and myotubes, with similar results found in the relative *SIX* mRNA levels (Fig. 6A-D). For both SIX1 and SIX2 overexpression, the low levels of *DUX4* and DUX4 target mRNAs in FSHD myoblasts were even lower in overexpressing lines (Fig. 6E). For the SIX1 overexpressing FSHD1 line, *DUX4* and DUX4 target *MBD3L2* were induced upon differentiation to an absolute level lower than that in the parental line. However, the relative increase in *DUX4* and target (*MBD3L2*) mRNA levels during myogenic differentiation were similar between parent FSHD1 and our SIX1 overexpressing FSHD1 lines when normalized to each respective myoblast cell line (Fig. 6E-H).

**Figure 6.**
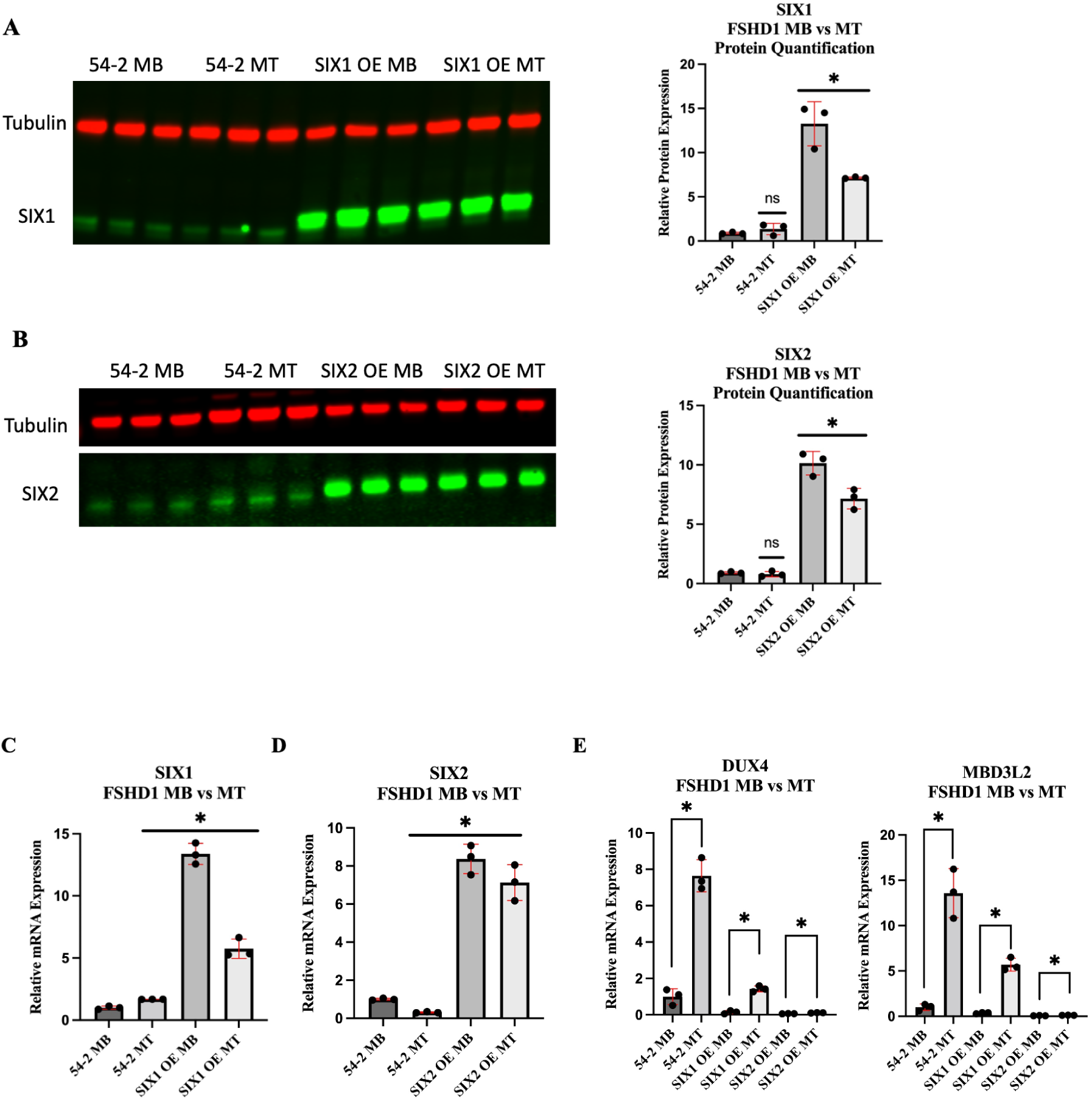
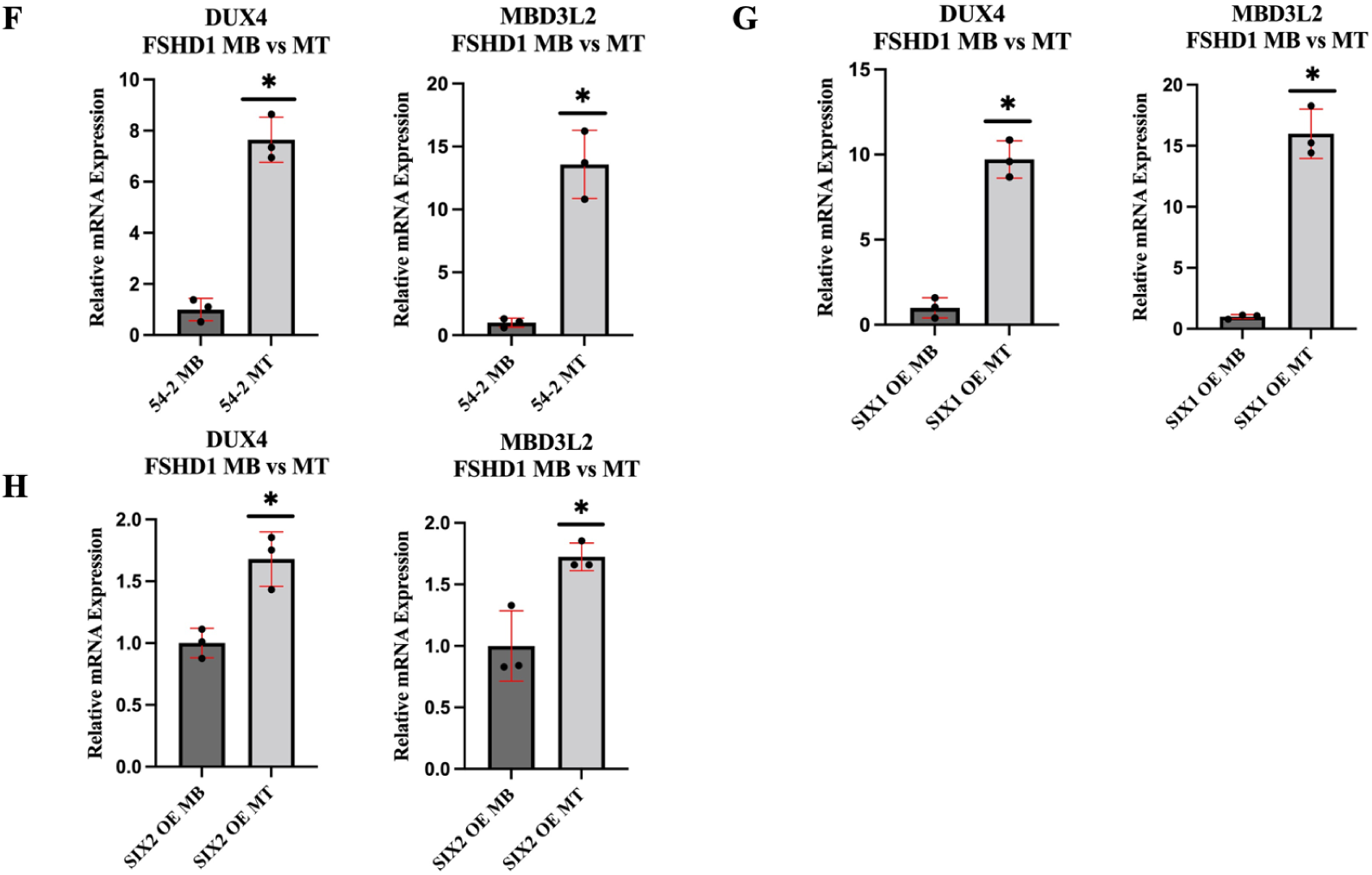
Overexpression of SIX1 and SIX2 are not limiting. Shown are FSHD1 (54-2) myoblasts lines overexpressing SIX1 and SIX2 produced by lentiviral vectors and compared to unmodified FSHD1 myoblasts to myotubes. (**A** and **B**) Western blot and quantification confirm overexpression of SIX1 (**A**) and SIX2 (**B**). Alpha-tubulin was used as the loading control. Relative mRNA levels for *SIX1* (**C**), *SIX2* (**D**), *DUX4* and DUX4 target*, MBD3L2* (**E**). From the data shown in (**E**), the relative induction of each cell line was normalized to its respective myoblast condition to demonstrate the induction of *DUX4* mRNA following differentiation (**F-H**). Each experimental condition (n=3) was normalized to a negative si- Control with the mean and standard deviation depicted. Asterisks demonstrate statistical significance between the control vs experimental group using an unpaired two tailed *t-*test (ns= not significance; *p<0.05).

The SIX2 overexpressing FSHD1 line behaved differently, where there was consistently a smaller increase in DUX4 target mRNA upon differentiation (Fig. 6E and Fig. 6H). Overall, these data demonstrate that increasing SIX1 or SIX2 protein levels does not increase *DUX4* or DUX4 target mRNA levels and therefore they are not limiting their abundance in turning on DUX4 expression.

### DUX4 suppresses *SIX* gene expression

Previous RNA-seq studies demonstrated that *SIX1*, *SIX2* and *SIX4* expression was decreased after forced DUX4 expression (Bosnakovski et al. 2018; Jagannathan et al. 2016). In light of the important role these factors play in driving *DUX4* expression, we hypothesized the existence of a negative feedback loop in which DUX4, induced in differentiating myotubes in a SIX transcription factor-dependent manner, would suppress SIX expression to limit its own transcription. Since DUX4 is induced in only a subset of nuclei during FSHD myogenic differentiation, this suppression would not be accurately assessed in analysis of RNA derived from the myocyte population. To overcome this limitation, we utilized immortalized non-FSHD human myoblasts (MB135) engineered with a doxycycline-inducible promoter driving *DUX4* so that the effects of DUX4 could be monitored in the population (Jagannathan et al. 2016). Induction of *DUX4* by treatment of MB135-iDUX myoblasts with doxycycline increased DUX4 target genes in a concentration and time-dependent manner with saturation of DUX4 target gene expression at 0.5-1.0 ug/ml doxycycline at 8 hours and 0.25-0.5 ug/ml at 24 hours (Fig. 7A-B). To determine the regulation of *SIX1*, *SIX2* and *SIX4* individually under high induction of DUX4, we treated the cells for 24 hours with a near saturating concentration of doxycycline and found *SIX1*, *SIX2*, and *SIX4* expression were each significantly decreased at a doxycycline concentration that models a burst of DUX4 expression, consistent with previous profiling studies (Fig. 7C-E) (Bosnakovski et al. 2018; Jagannathan et al. 2016). These results suggest that high expression of DUX4 suppresses *SIX1*, *SIX2*, and *SIX4* in a negative feedback loop.

**Figure 7.**
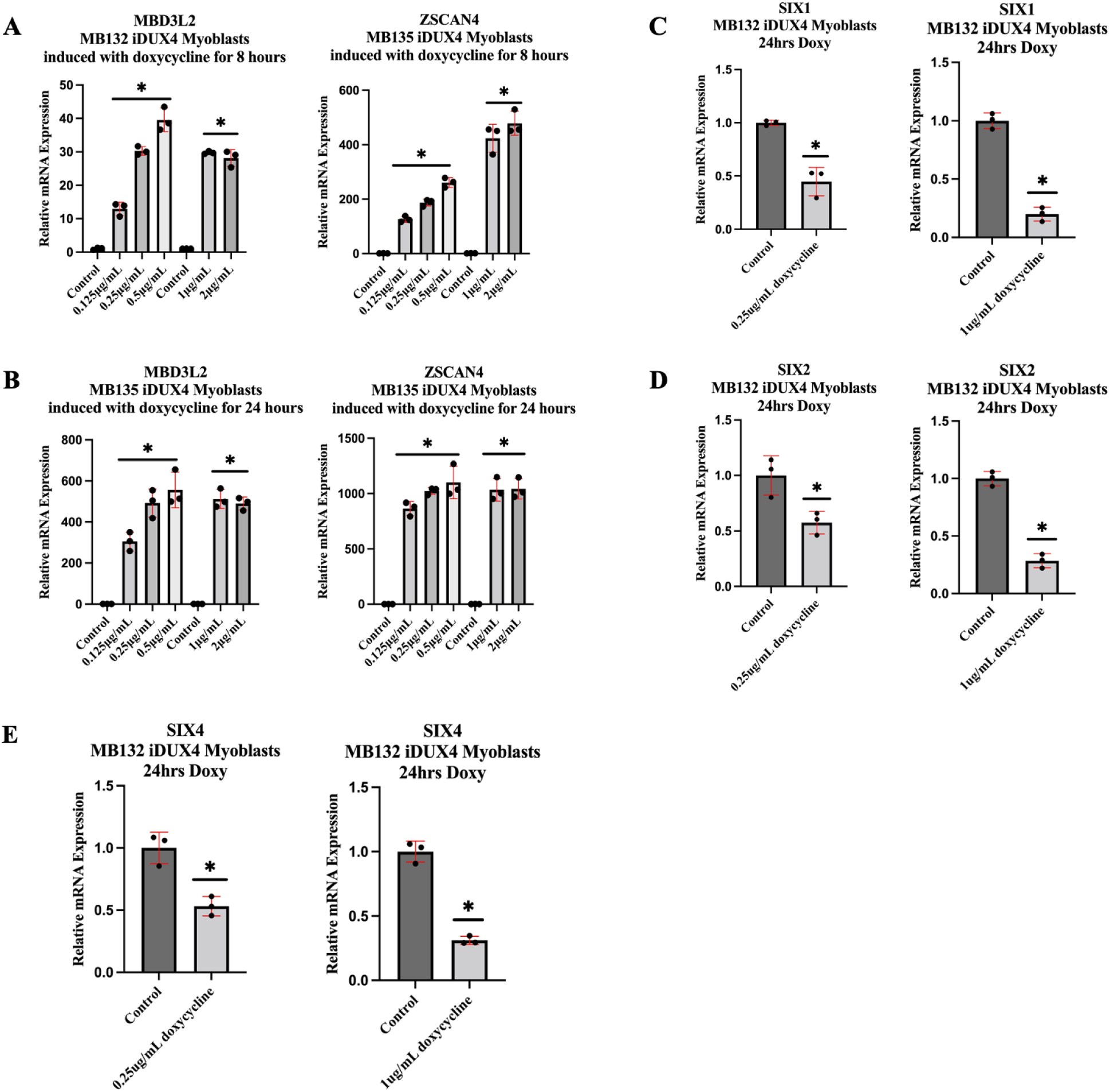
Forced DUX4 expression downregulates SIX1, SIX2 and SIX4. Stable non-FSHD (MB135) doxycycline-inducible DUX4 myoblasts were induced with various concentrations of doxycycline (0.125, 0.25, 0.5, 1, and 2ug/mL) for 8 hours (**A**) and 24 hours (**B**). Shown are the relative mRNA levels for DUX4 targets *MBD3L2* and *ZSCAN4* (**A** and **B**), *SIX1* (**C**), *SIX2* (**D**) and SIX4 (**E**) at the indicated concentrations of doxycycline. Each experimental condition (n=3) was normalized to a negative si-Control with the mean and standard deviation depicted. Asterisks demonstrate statistical significance between the control vs experimental group using an unpaired two tailed *t-*test (ns= not significance; *p<0.05).

## Discussion

The transcription factor DUX4 plays a critical role in early embryonic development, when it is involved in zygotic genome activation to orchestrate developmental gene expression before being silenced in most somatic tissues (Mocciaro et al. 2021; Statland and Tawil 2014; van der Maarel, Tawil, and Tapscott 2011; Brouwer et al. 1994). Its inappropriate expression in adult skeletal muscle causes the progressive myopathic disease FSHD (Lemmers et al. 2010). Additionally, DUX4 expression is activated in a subset of cancers, where it promotes immune evasion, resistance to checkpoint blockade and immunotherapy failure in metastatic disease (Chew et al. 2019; Pineda and Bradley 2023). Consequently, there is immense interest in DUX4 as a therapeutic target and understanding the factors that promote its expression is critical to therapeutics development (Karpukhina et al. 2021; Sidlauskaite et al. 2020; Tihaya et al. 2023).

The hallmark of FSHD patient-derived muscle cells in tissue culture is myogenic differentiation-dependent transcriptional bursts of *DUX4* in rare nuclei (Block et al. 2013; Rickard, Petek, and Miller 2015; Snider et al. 2010; Tassin et al. 2013). In FSHD, loss of repeat-mediated epigenetic repression at D4Z4 macrosatellite repeats is observed with specific decreases in repressive epigenetic marks including DNA methylation, histone 3 lysine 9 trimethylation (H3K9me3) and H3K27me3 (Tawil, van der Maarel, and Tapscott 2014); however, there is little detail on the complex transcriptional activation events that ultimately produce *DUX4* transcripts. While the restriction of these bursts to rare nuclei remains to be understood, it is presumed that myogenic differentiation signals drive *DUX4* expression in those nuclei that have escaped epigenetic silencing, and indeed myogenic enhancers upstream of D4Z4 repeats have been described, although no specific transcription factors were identified that directly contribute (Himeda et al. 2014; Sidlauskaite et al. 2020; Tawil, van der Maarel, and Tapscott 2014; Tihaya et al. 2023). A well-established myogenic differentiation signal is p38 MAPK activation, which we and others have shown positively regulates transcription of *DUX4*, both *in vivo* and *in vitro*, supporting p38 as a drug target in FSHD (Oliva et al. 2019; Rojas et al. 2020). Yet, the dynamic interactions between p38 MAPK and the factors required for *DUX4* regulation remain to be elucidated. Modulation of other signaling pathways and effectors also regulate *DUX4* including Wnt/b-catenin activation, b2-adrenergic agonism/protein kinase A activation, bromodomain and extra-terminal domain (BET) inhibition, casein kinase 1 d/e inhibition and other kinase pathways, yet the intersection of these pathways with *DUX4* transcription has yet to be defined (Block et al. 2013; Campbell et al. 2017; Cruz et al. 2018; De Maeyer J 2022; Rickard 2023).

In this study, we demonstrate that SIX transcription factors drive myogenic differentiation-dependent bursts of DUX4 in patient-derived muscle cells. We found, using combined siRNA knockdown of SIX1, SIX2 and SIX4, that *DUX4* expression was nearly completely suppressed in FSHD1 (54-2 and 16- ABIC) and FSHD2 (MB200) myotubes. Combinations containing *SIX2* siRNA were the most effective at reducing *DUX4* mRNA levels, highlighting the particular importance of SIX2 in driving *DUX4* transcription. Suppression of *DUX4* was context-dependent, occurring only in differentiating cells in which *DUX4* expression bursts are induced. Knockdown of SIX1, SIX2 and SIX4 in myoblasts failed to affect the low-level expression of *DUX4* in undifferentiated FSHD myoblasts. We demonstrated that SIX1 is transcriptionally active in myoblasts, since knockdown of *SIX1* in FSHD and non-FSHD myoblasts dramatically reduced RNA levels for *PGK1*, a glycolytic gene known to be regulated by SIX1 (Li et al. 2018). Since SIX protein levels are not induced during differentiation, this data indicates that SIX protein activity in promoting *DUX4* expression is likely dependent on myogenic signaling. As mentioned above, one such aspect of myogenic signaling is p38 MAPK activation, which orchestrates genome-wide transcriptional changes (Zetser, Gredinger, and Bengal 1999; Segales et al. 2016).

Although SIX factors are not known to be regulated by phosphorylation, it is possible that p38 modulates their interaction with necessary cofactors. Another aspect of myogenic signaling may be the availability of other active transcription factors acting synergistically with SIX proteins, a common theme for SIX transcription factor activities in driving different stages of muscle development (Maire et al. 2020). An alternative interpretation is that the chromatin environment is changed during differentiation to allow SIX transcription factors access to DUX4 regulatory regions. The potential regulation of SIX transcription factors by myogenic signaling pathways such as p38 warrants further investigation.

Due to the potential involvement of p38 modulating SIX1, SIX2, or SIX4 activities through phosphorylation of known cofactors, we investigated the potential regulation of *DUX4* transcription by the SIX coactivator, EYA. *In vitro*, p38a/b was found to phosphorylate and activate EYA (Hsiao et al. 2001); therefore, we tested the involvement of EYA (EYA1, EYA3, and EYA4) in the regulation of *DUX4*. Using a combined siRNA knockdown of EYA1, EYA3, and EYA4, we found that *DUX4* expression was partially suppressed in FSHD1 myotubes, demonstrating an involvement in DUX4 regulation. Individual and combined knockdowns containing EYA3 had the most pronounced effect, consistent with EYA3 expression being most widespread (Soni, Roychoudhury, and Hegde 2021).

Transcriptional regulation mediated by the SIX-EYA complex has been widely explored. Cytoplasmic EYA proteins, when complexed with SIX, are translocated to the nucleus where together they activate target gene expression (Soni, Roychoudhury, and Hegde 2021; Maire et al. 2020). Interestingly, EYA has been found to interact directly with chromatin remodeling complex ATPases to allow for the SIX-EYA complex to promote DNA accessibility and initiate transcriptional activation (Maire et al. 2020; Ahmed, Xu, and Xu 2012). Given the involvement and importance of p38 MAPK’s recruitment of chromatin remodeling complexes and the proposed activation of EYA by p38 phosphorylation, this potential mechanism needs to be further explored (Simone et al. 2004; Hsiao et al. 2001). Alternatively, SIX2 has been shown to directly recruit the SWI/SNF chromatin remodeling complex independent of EYA (Gao et al. 2021), and is known to activate transcription in the absence of EYA cofactors (Blevins et al. 2015). Our data suggests that SIX proteins utilize both EYA-dependent and EYA-independent mechanisms to drive DUX4 expression.

Important for making inferences about *DUX4* regulation, we demonstrated that knockdown of SIX genes does not inhibit myogenic differentiation. Multinucleated myotubes formed similarly between control cultures and combined SIX1, SIX2, and SIX4 knockdown cultures in two FSHD1 (54-2 and 16-ABIC) and one FSHD2 (MB200) patient-derived cell lines. Markers of differentiation such as *MYOD1*, *MYOG* and *MYH1* were unaffected. One late differentiation marker, creatine kinase, M-type (*CKM*), was significantly decreased by combined SIX knockdown; however, *CKM* is known to be directly regulated by SIX transcription factors (Himeda et al. 2004). RNA levels for *MYH3* and *MYH8*, genes encoding developmental and regenerative myosin heavy chains, were both decreased by combined knockdown.

Interestingly, we found differential effects on genes involved in myofiber type specification. While slow myofiber type gene, *MYH*7, was dramatically decreased, fast fiber type, *MYH2,* was increased by combined SIX1/2/4 knockdown. SIX transcription factors have been found to accumulate preferentially in the nuclei of Type 2 fast-fiber skeletal muscles, the fibers preferentially affected with FSHD (Sakakibara et al. 2016; Maire et al. 2020). It will be important to investigate the potential connection between muscle fiber types expressing SIX family members and those that are selectively lost in degenerating FSHD muscle. For example, FSHD affects fast-twitch oxidative (MYH2; Type 2A) and glycolytic muscle fibers (MYH1, Type 2X), reducing maximum force capacity in select muscle groups that are affected (Celegato et al. 2006; Lassche et al. 2013). Nonetheless, since combined SIX knockdowns did not prevent myogenic differentiation, we were able to demonstrate their specific critical role in *DUX4* induction during differentiation.

The restricted and sporadic nature of DUX4 being expressed in only a subset of nuclei is a poorly understood phenomenon of FSHD. *In vitro*, DUX4 is expressed at relatively low levels in undifferentiated myoblasts; however, following differentiation, there is a burst of DUX4 in rare sentinel nuclei (Rickard, Petek, and Miller 2015). In this study, we demonstrated the requirement of SIX1/2/4 to induce the transcriptional activation of *DUX4* expression. Because DUX4 could be regulated by several factors, we wanted to determine if its restricted expression pattern was due to the absence or presence of necessary transcription factors, like SIX1 and SIX2. We found that although SIX1 and SIX2 are necessary for DUX4 expression, they are present in every myonuclei in our FSHD cultures and thus their expression pattern does not explain the restricted expression of DUX4 to a subset of nuclei. These data support the idea that SIX transcription factors require some other rare myogenic signaling event or cumulative epigenetic injury to escape silencing.

Since we found SIX1 and SIX2 to be critical drivers of *DUX4* transcription, we tested whether their overexpression would further induce *DUX4* mRNA levels. We found that overexpression of either SIX1 or SIX2 resulted in reduced DUX4 target mRNA levels compared to the parental FSHD1 cell line. From this observation, we determined that SIX1 and SIX2 are not limiting in their abundance to turn on DUX4 expression, although the high level of overexpressed transcription factors may have simply exceeded the cells’ capacity in terms of available cofactors. A more precise titration of SIX levels may be necessary to see subtle differences. Additionally, we investigated the potential for feedback regulation between DUX4 and SIX1, SIX2 and SIX4. We used a doxycycline-inducible DUX4 cell model that would allow us to express DUX4 in every cell nuclei and study the regulation between DUX4 and the SIX genes. We found that the high induction of DUX4 expression by doxycycline resulted in the downregulation of *SIX1*, *SIX2* and *SIX4*, suggesting negative feedback regulation. It has been demonstrated that ectopic DUX4 expression can lead to several feedback and feedforward mechanisms contributing to its toxic persistent expression in the disease state (Jagannathan 2022).

Conversely, in embryonic development, DUX4 is expressed in a discreet window, inducing zygotic genome activation (ZGA) before being silenced as development proceeds (De Iaco et al. 2017; Hendrickson et al. 2017). It will be important to identify drivers of DUX4 during ZGA, the mechanisms of suppression, and similarities and differences with FSHD muscle cells.

Where do SIX factors bind to exert their effect on *DUX4* transcription, and do they act directly? Himeda *et al*. described myogenic enhancers upstream of the 4q D4Z4 repeats that activate *DUX4* expression in FSHD skeletal myocytes. In particular, *DUX4* myogenic enhancer 2 (DME2) exhibits a strong muscle-specific interaction with the *DUX4* promoter and *in silico* analysis of the sequence revealed the presence of a consensus binding motif for SIX1/4 (MEF3) as well as other regulators of muscle gene expression (Himeda et al. 2014). This suggests the possibility that SIX proteins directly activate *DUX4* transcription through DME2. It is not clear if SIX1/2/4 are functionally acting at the same site or if they participate in separate steps of *DUX4* transcription by binding at multiple locations and/or during sequential steps within a transcription cascade. It will be important to experimentally determine genomic binding regions for each SIX family member to understand their individual roles in promoting *DUX4* activation.

SIX transcription factors are known master regulators during the development of the head, sensory organs, kidney and skeletal muscle through their control of progenitor cell populations and differentiation mechanisms (Meurer et al. 2021; Maire et al. 2020; Davis and Rebay 2017; Inoue et al. 2023). Interestingly, *SIX* genes are expressed developmentally in the pre-placodal ectoderm and later in sensory cranial placodes, precursors of sensory organs of the head (Moody and LaMantia 2015; Park and Saint-Jeannet 2010). Recently, it has been shown that mutations in SMCHD1, most commonly associated with FSHD2, are also responsible for the congenital defect arrhinia, complete lack of the external nose (Gordon et al. 2017; Shaw et al. 2017). Inoue *et al*. demonstrated a potential connection with DUX4 expression by showing that defects in SMCHD1 can allow DUX4 expression during cranial placode differentiation, causing DUX4 toxicity and cell death, providing a plausible link between SMCHD1 mutations and developmental defects such as arrhinia (Inoue et al. 2023). It is tempting to speculate that SIX transcription factors are a common link to cells capable of expressing DUX4 and that other co-morbidities in FSHD, such as high frequency hearing loss, may be due to a similar toxicity in which DUX4 is expressed during development of otic placodes. Understanding the nature of the regulation of DUX4 expression by SIX transcription factors will spur a deeper understanding of the pathogenesis of FSHD, comorbidities potentially linked to disease severity in FSHD, and cancer. Further investigation will be needed to identify other transcriptional and epigenetic factors regulating DUX4 expression to develop a mechanistic understanding of pathogenic DUX4 expression.

## Materials and Methods

### Experimental Design

The objective of this study was to identify novel transcriptional regulators of DUX4 expression to expand our understanding of the factors promoting FSHD pathogenesis. To do this, we utilized immortalized patient derived FSHD cells to study the effects on *DUX4* transcription by siRNA knockdown of SIX transcription factors (SIX1, SIX2 and SIX4). Early and late differentiation markers were analyzed by qPCR analysis to monitor any differential changes following siRNA knockdown. Immunofluorescence was used to look for correlations between SIX proteins and the restricted expression of DUX4 in rare myonuclei. Lentiviral vectors were utilized to study overexpression of SIX1 and SIX2, due to their most pronounced role in promoting *DUX4* expression.

Non-FSHD myoblasts with a doxycycline-inducible *DUX4* transgene were utilized to study the regulation of DUX4 expression on *SIX* gene transcription. Our data provides strong evidence of the involvement of SIX genes in the regulation of DUX4.

### Cell Culture System

FSHD1 (54-2; 3 D4Z4 repeat units), FSHD1 (16-ABIC; 7 D4Z4 repeat units), and FSHD2 (MB200) patient-derived immortalized myoblasts and non-DUX4 expressing cells (54-6; 13 D4Z4 repeat units) were grown in Ham’s F-10 Nutrient Mix (Gibco, Waltham, MA) with 20% FBS (Corning, Corning, NY), 100 U/100µg penicillin-streptomycin (Gibco), 10ng/mL of recombinant human FGF (Promega Corporation, Madison, WI), and 1µM dexamethasone (Sigma-Aldrich, Saint Louis, MO). To induce differentiation to form multi-nucleated myotubes, we used Dulbecco’s modified Eagle’s medium/F-12 mix (Gibco) with 100 U/100µg penicillin-streptomycin, and 10µg/mL knockout serum (Gibco) and differentiated our cells for 40-48 hours. For our inducible DUX4 cell system (iDUX4), we used codon-altered MB135 cells with a doxycycline-inducible promoter gifted by Dr. Stephen Tapscott. MB135 iDUX4 cells were supplemented with 3ug/mL puromycin (Thermo Fischer Scientific, Waltham, MA) for selection. DUX4 was induced with doxycycline (Thermo Fischer Scientific) for 8 and 24 hours.

### Transfections of small interfering RNA (siRNA)

The Silencer Select siRNAs for human SIX1, SIX2, and SIX4 were purchased from Thermo Fischer Scientific (Waltham, MA). We plated the cells at 1x10^5^ in a 12-well plate and transfected the cells hours later once the cells began adhering to the plate. To transfect the siRNAs in our proliferating myoblasts, we optimized the amount of Opti-MEM Reduced Serum Medium (Gibco, Waltham, MA) and Lipofectamine RNAiMAX (Thermo Fisher Scientific, Waltham, MA) needed for a 12-well plate, and transfected 3µL of 10pmol of the target gene of interest and compared a non-targeting siRNA control, with each condition being carried out in triplicate. 72 hours after transfecting the cells, myoblasts were harvested using lysis buffer and the RNA was isolated using the E.Z.N.A. Total RNA Kit 1 (Omega BioTek, Norcross, GA). In conditions where we wanted to observe the effects in myotubes, 48 hours after the transfection to ensure depletion of the target gene, we switched the growth media to a differentiation media to allow for myotubes to form. We differentiated our cells for 40-48 hours prior to harvesting and isolating the RNA for qPCR analysis.

### siRNA Sequences

The siRNAs were all purchased from Thermo Fisher Scientific (Waltham, MA). Silencer Select Negative Control No.1 (cat. 4390843) was used throughout all experiments. SIX1 (s12874, sense: AGAACGAGAGCGUACUCAAtt,antisense: UUGAGUACGCUCUCGUUCUtg); SIX2 (s21094, sense: GGGAAUAAAUUAUACACCAtt, antisense: UGGUGUAUAAUUUAUUCCCtt); SIX4 (s224246, sense: GGUUGAUACUGUCUGUGAAtt, antisense: UUCACAGACAGUAUCAACCat).

### Total RNA Extraction and qPCR

Total RNA was extracted using spin-columns from the E.Z.N.A Total RNA Kit purchased from Omega BioTek according to the kit instructions (Norcross, GA). Quantitative real-time polymerase chain reaction (qRT-PCR) was used to analyze to relative expression levels using Quant Studio 5 from Applied Biosystems by Thermo Fischer Scientific (Foster City, CA).

### TaqMan Assay for qPCR

All TaqMan primer probe sets were purchased from Thermo Fisher Scientific (Waltham, MA) and compared to an endogenous control, RPL30 (Hs00265497_m1). Human DUX4 and DUX4 targets were used: DUX4 (Hs07287098_g1); MBD3L2 (Hs00544743_m1); LEUTX (Hs01028718_m1); and ZSCAN4 (Hs00537549_m1). Human SIX genes were used: SIX1 (Hs00195590_m1); SIX2 (Hs00232731_m1); SIX4 (Hs00213614_m1). To look at markers of differentiation, MYOG (Hs01072232_m1); MYOD (Hs00159528_m1); MYH1 (Hs00428600_m1); MYH2 (Hs00430042_m1); MYH3 (Hs01074230_m1); MYH4 (Hs00757977_m1); MYH7 (Hs01110632_m1); MYH8 (Hs00267293_m1); CKM (Hs00176490) were used. SIX1 and SIX2 target genes were used: PGK1 (Hs00943178_g1); SLC4A7 (Hs00186192_m1); EYA1 (Hs00166804_m1). For each assay, TaqMan Fast Virus 1-Step Master Mix (Thermo Fisher Scientific, Walham, MA) or TaqMan Fast Advanced Virus Master Mix (Thermo Fisher Scientific) were used to carry out qPCR.

### DUX4 Assay

To determine relative *DUX4* expression, a two-step process was used in which cDNA was first synthesized with oligo dT priming using the ProtoScript II First-Strand cDNA Synthesis kit (New England BioLabs, Ipswich, MA). To denature the template RNA, 6µL of total RNA of interest with 2µL of Oligo dT and incubated for 5 minutes at 65°C. Then 10uL of the Reaction Mix (2X) and 2uL of the Enzyme Mix (2X) was mixed with the RNA. Samples were incubated for 1 hour at 42°C and then enzyme was inactivated for 5 minutes at 80°C. cDNA was used in the standard qPCR protocol.

### Western Blot

Cells treated by siRNA knockdown were collected by adding direct SDS lysis buffer to each well. The samples were then collected and boiled for 10 minutes. After boiling, the samples were run on a NuPAGE 4-12% Bis-Tris polyacrylamide gel (Thermo Fisher Scientific, Waltham, MA) using an SDS Running buffer (Thermo Fisher Scientific) and transferred to an Immobilon-FL polyvinylidene difluoride transfer membrane (Millipore, Burlington, MA) using the NuPAGE transfer buffer (Life Technologies, Carlsbad, CA) with 10% methanol. After the membrane transfer, the membrane was left to dry overnight. The next day, the membrane was activated by soaking in methanol and 1X TBS before blocking in Odyssey TBS Blocking buffer (Li-Cor, Lincoln, NE) for 1 hour. After blocking, the primary antibody of interest and an appropriate α/β-Tubulin (loading control) were added to the membrane at a concentration of 1:1000 in Odyssey TBS Blocking buffer (Li-Cor) and 0.2% Tween 20 overnight at 4°C. The next day the membrane was washed using 1X-TBST (0.1% Tween 20) before adding a near-infrared fluorescent secondary antibody (Li-Cor) diluted at 1:15,000 in the Odyssey TBS Blocking buffer (Li-Cor) with 0.2% Tween 20 and 0.01% SDS. The membrane was incubated in the secondary antibody for 1 hour at room temperature. After a series of washes with 1X TBST (0.1% Tween 20) and a rinse of 1X TBS to remove all the Tween 20, the Li-Cor Odyssey CLx was used to image the blot. Knockdown analysis was carried out on Empiria Studio software to determine the percentage of selective protein knockdown.

### Primary and Secondary Antibodies

The primary antibodies that were used for Western Blotting were α- Tubulin Mouse (Li-Cor; 926-42213), β-Tubulin Rabbit (Li-Cor; 926-42211), SIX1 Rabbit mAb (Cell Signaling Technologies, Danvers, MA; 12891S), SIX2 Rabbit pAb (Novus, Centennial, CO; NBP2- 54917), and SIX4 Rabbit pAb (Abcam, Cambridge, UK; ab176713). The secondary antibodies that were used were Goat anti-Rabbit IRDye 680RD (Li-Cor; 926-68071), Goat anti-Mouse IRDye 680RD (Li-Cor; 926-68670), Goat anti-Mouse IRDye 800CW (Li-Cor; 926-32210), and Goat anti-Rabbit IRDye 800CW (Li-Cor; 926-32211).

### Lentivirus production

Lentivirus was produced in HEK293T cells, using standard triple transfection protocols. Briefly, HEK293T were co-transfected with lenti-vector pMDLg/pRRE (Addgene no. 12251), pRSV-Rev (Addgene no. 12253), pMD2.G (Addgene no. 12259) and respective individual plasmids listed above using OptiMEM and Lipofectamine 3000 reagent. After 48 and 72 hours, the supernatant was collected and filtered through a 0.45 μm strainer. It was followed by lentivirus concentration, after which the virus titer was calculated, and stored at -80 °C. Lentiviral vectors expressed SIX1 (plasmid #120476) and SIX2 (plasmid #120477) with control under an EF1α promoter (Addgene, Watertown, MA).

### Generation of stably expressing SIX FSHD cells

FSHD1 (54-2) cells were transfected with SIX1 and SIX2 lentivirus (MOI < 0.3) in the presence of 2 µg/ml polybrene followed by selection with hygromycin (500 ng/ml) for 7 days. Western blot analysis was used to confirm the overexpression of SIX protein expression.

### Immunofluorescence

Cells were fixed using 4% paraformaldehyde (Thermo Fischer Scientific) for 20 minutes at room temperature, permeabilized with 2% Triton for 15 minutes, washed with 1X PBS (Corning), blocked with 5% donkey/or goat serum (Millipore) and 1% BSA (Thermo Fischer Scientific). Primary antibodies were incubated at 1:300 at 4°C overnight. The primary antibodies used were SIX1 Rabbit mAb (Cell Signaling Technologies, Danvers, MA; 12891S) and SIX2 Rabbit pAb (Novus, Centennial, CO; NBP2-54917). The next day the cells were washed with 1X PBS and incubated with a secondary antibody (1:300) at room temperature for 1 hour. The secondary antibody used included Alexa Fluor 488 donkey anti-rabbit (Abcam; A21206), Alexa Fluor 488 goat anti-rabbit (Abcam; AB150077), Alexa Fluor 594 goat anti-mouse (Thermo Fischer Scientific; A11005) and DAPI (Thermo Fischer Scientific; 62248). Cells were then mounted and imaged using a Keyence Imaging Microscope under 40X magnification. For fusion index, the total nuclei and nuclei detected in myotubes were manually calculated. Clusters of nuclei (>2) per myotube was the standard in calculating the fusion index.

### Statistical Analysis

All figures were generated using GraphPad Prism 9. Statistical analysis was conducted using GraphPad Prism 9. All samples tested were normalized to the non-targeting control group. An unpaired two-tailed *t-test* was used to determine statistical significance by their individual p-values. Samples were considered significant with a p value <0.05.

## Acknowledgments

We would like to thank Joel Eissenberg for critical review of the manuscript. We thank Dr. Stephen Tapscott for providing MB135 iDUX4 cells.

## Funding

Muscular Dystrophy Association grant 576054 (FMS)

Friends of FSH Research (RV)

The Chris Carrino Foundation (RV)

The FSHD Society (RV)

## Author contributions

Conceptualization: AF, FS, JO, RV

Methodology: AF, FS, JO, RV

Investigation: AF, FS, JO, RV

Supervision: FS

Writing—original draft: AF, FS

Writing—review & editing: AF, FS, JO, RV

**Competing interests**: Authors declare that they have no competing interests.

## Data and materials availability

All data and materials will be available to any researcher for purposes of reproducing or extending the analyses. Materials transfer agreements (MTAs) with separate institutions are required for cell lines (see Methods). All data are available in the main text or the supplementary materials.

## Supplementary Materials

**Fig. S1.**
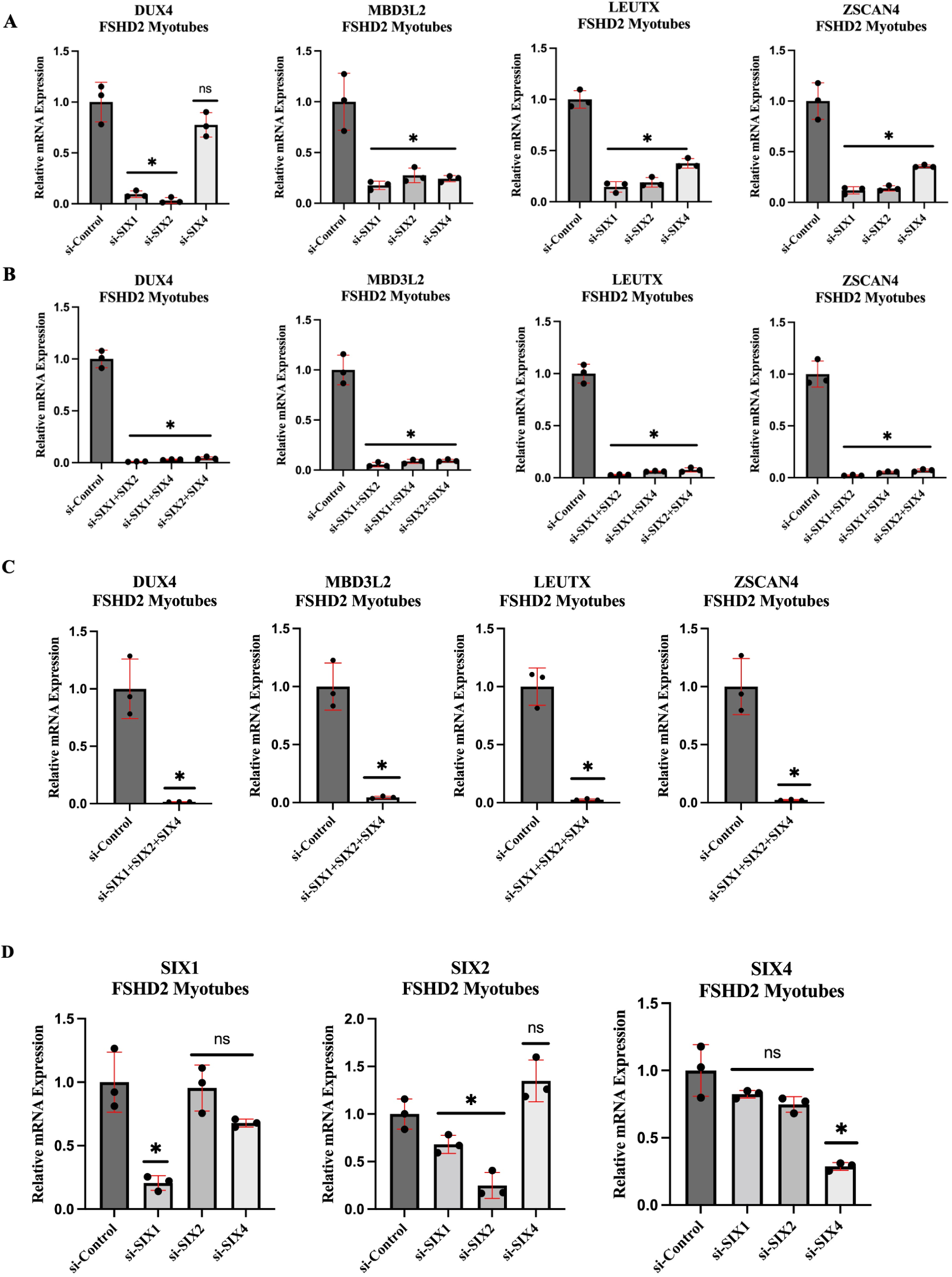
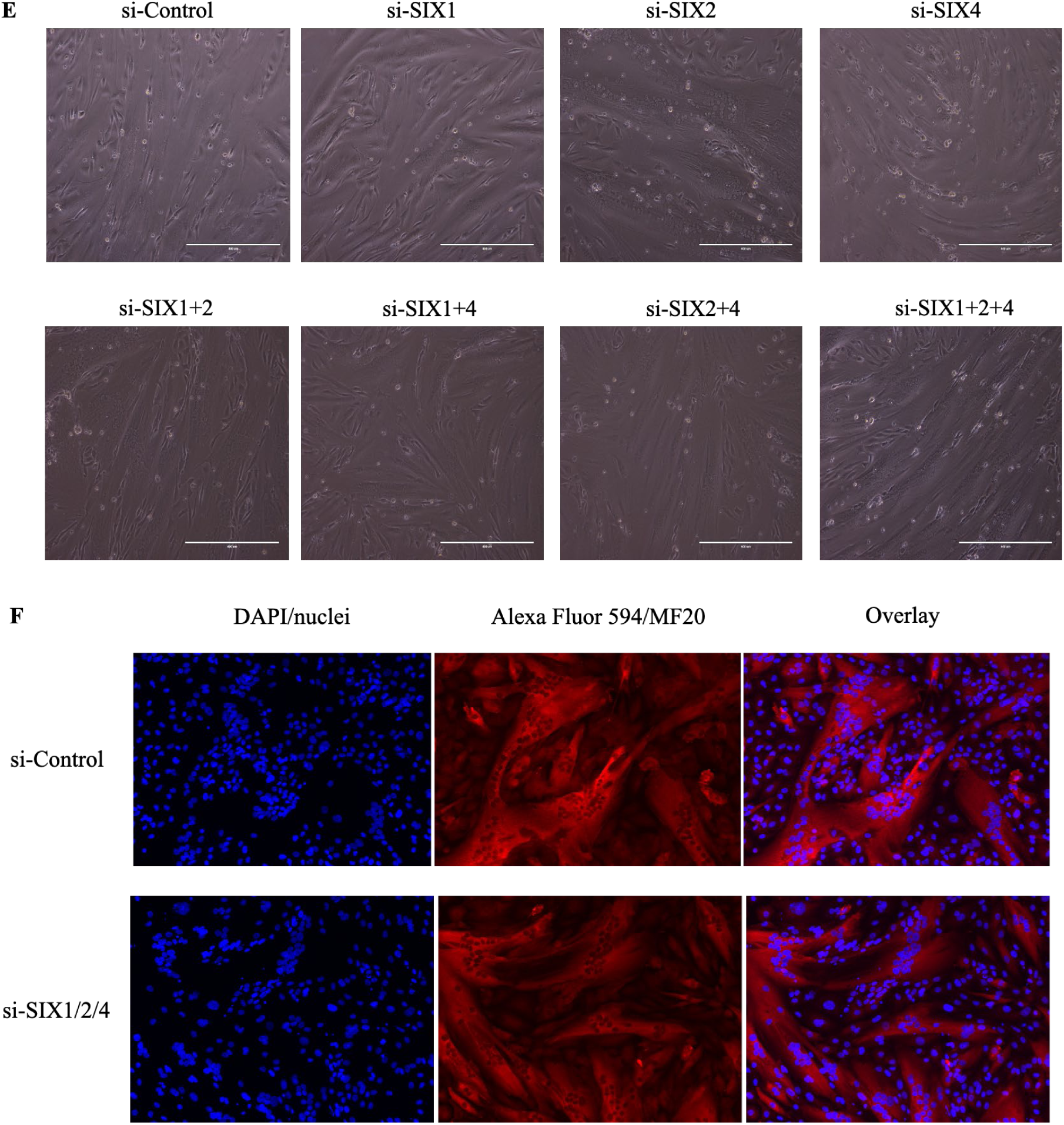
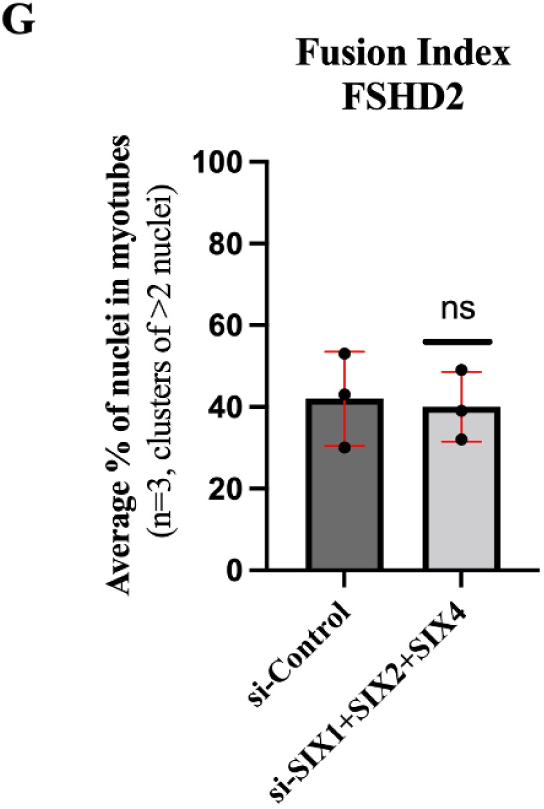
SIX1, SIX2, and SIX4 siRNA knockdown in FSHD2 differentiating muscle cells. FSHD2 (MB200) myoblasts were transfected with SIX1, SIX2, SIX4 siRNAs individually (**A**) and combinatorially (**B-C**) and differentiated to form multinucleated myotubes. Relative mRNA levels are shown for *DUX4* and its target genes (*MBD3L2*, *LEUTX* and *ZSCAN4)* (**A-C**) and each *SIX* paralog (**D**). (**E**) Brightfield images showing the morphology of multinucleated myotubes for each experimental condition. (**F**) Immunofluorescence staining for myosin heavy chain (MF20, red) and counterstained nuclei (blue) for the combined knockdown. (**G**) Fusion index was calculated (n=3) for clusters of myotubes (>2) for conditions shown in **F**. Each experimental condition (n=3) was normalized to a negative si-Control with the mean and standard deviation depicted. Asterisks demonstrate statistical significance between the control vs experimental group using an unpaired two tailed *t-*test (ns= not significance; *p<0.05).

**Fig. S2.**
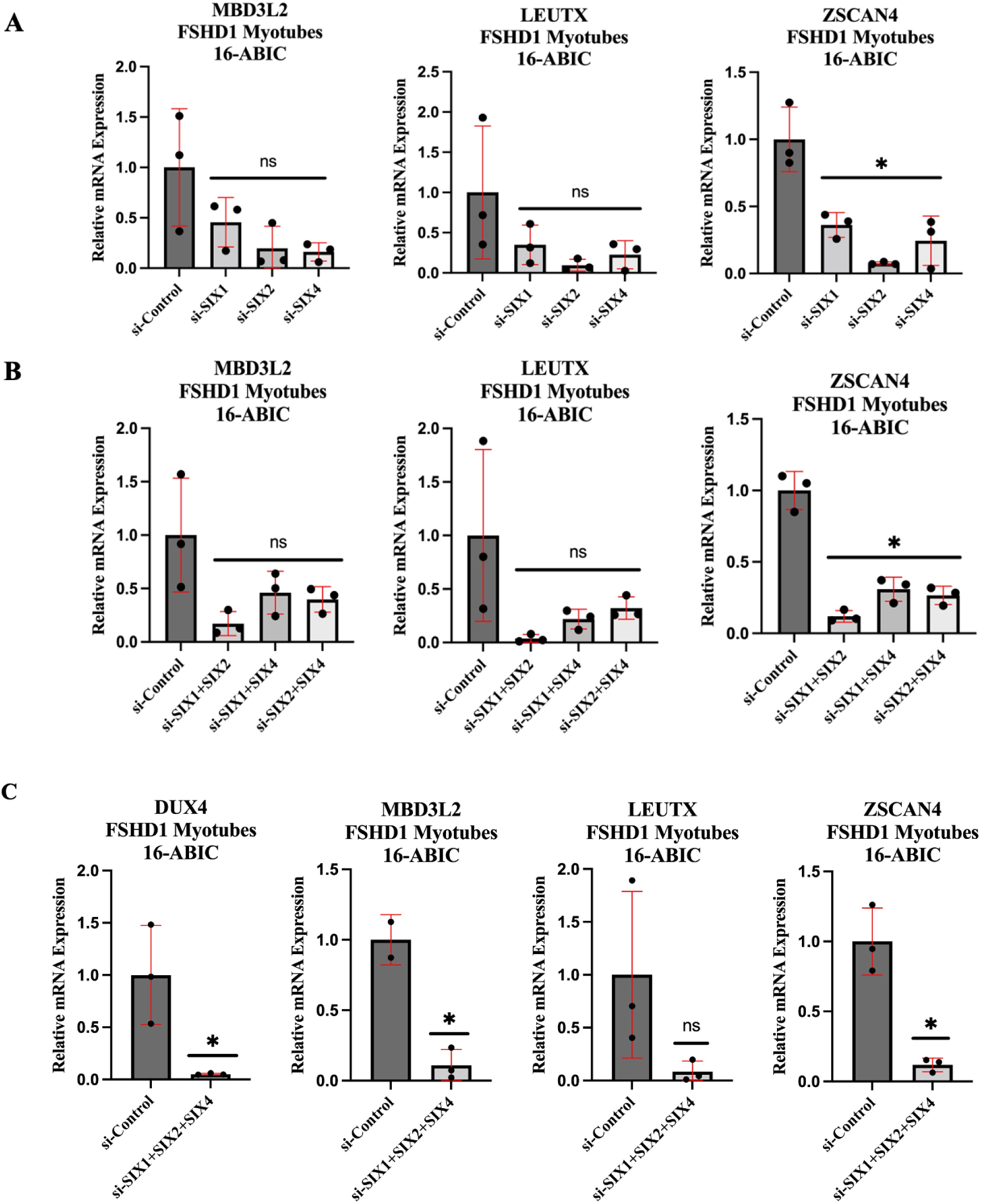
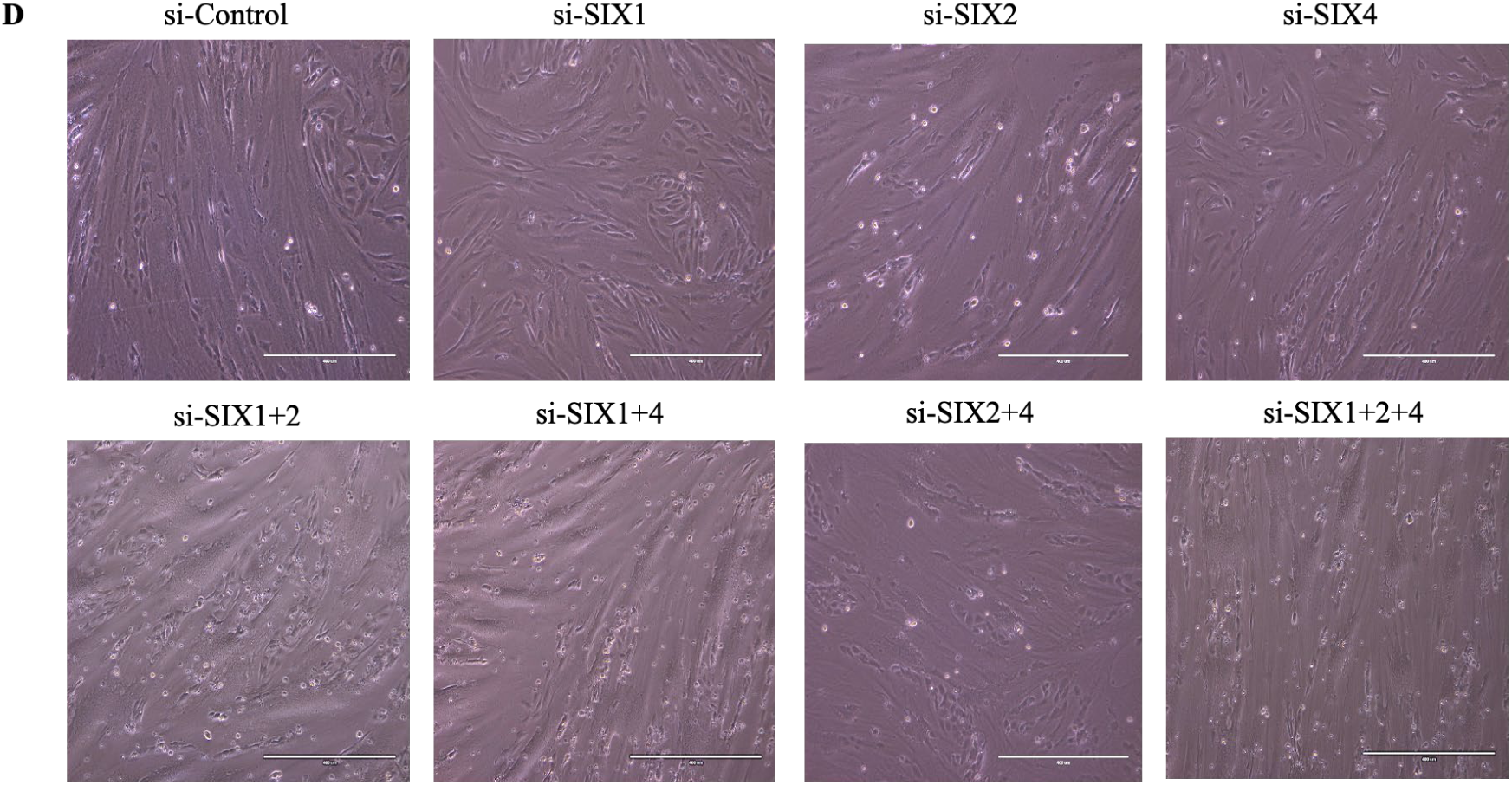
SIX1, SIX2, and SIX4 knockdown in FSHD1 (16-ABIC) myotubes. FSHD1 (16-ABICs) myoblasts were transfected with SIX1, SIX2, SIX4 siRNAs individually (**A**) and various combinations (**B-C**) similarly to Figure 1 and Fig. S1. mRNA levels for DUX4 target genes (*MBD3L2*, *LEUTX* and *ZSCAN4*) (**A-C**) and *DUX4* (**C**). (**D**) Morphology of the multinucleated myotubes was examined for each experimental condition. Each experimental condition (n=3) was normalized to a negative si-Control with the mean and standard deviation depicted. Asterisks demonstrate statistical significance between the control vs experimental group using an unpaired two tailed *t-*test (ns= not significance; *p<0.05).

**Fig. S3.**
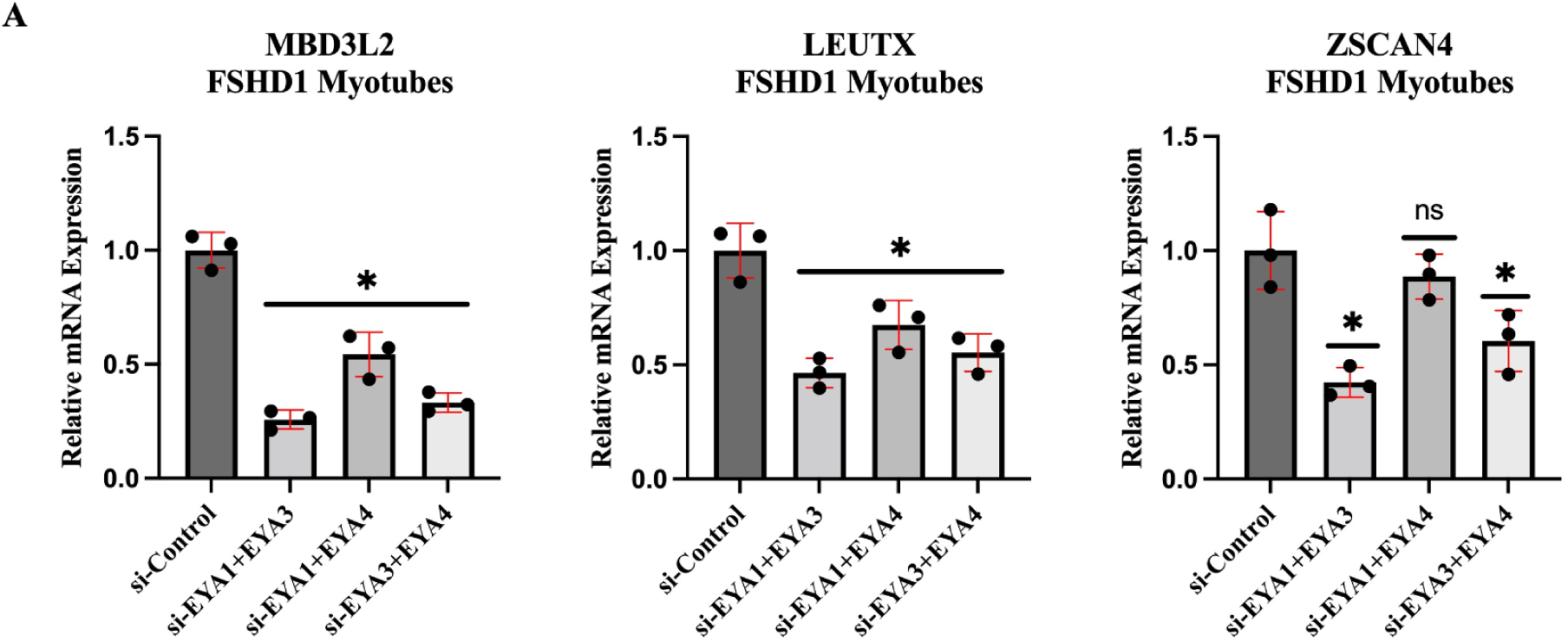
Dual combinations of EYA1/3/4 knockdown decreases DUX4 target expression levels. FSHD1 (54-2) myoblasts were transfected with dual combinations of highly expressed EYAs, EYA1/3/4, prior to inducing differentiation. (**A**) Shown are the relative mRNA levels of DUX4 targets (*MBD3L2*, *LEUTX* and *ZSCAN4*). Each experimental condition (n=3) was normalized to a negative si-Control with the mean and standard deviation depicted. Asterisks demonstrate statistical significance between the control vs experimental group using an unpaired two tailed *t-*test (ns= not significance; *p<0.05).

**Fig. S4.**
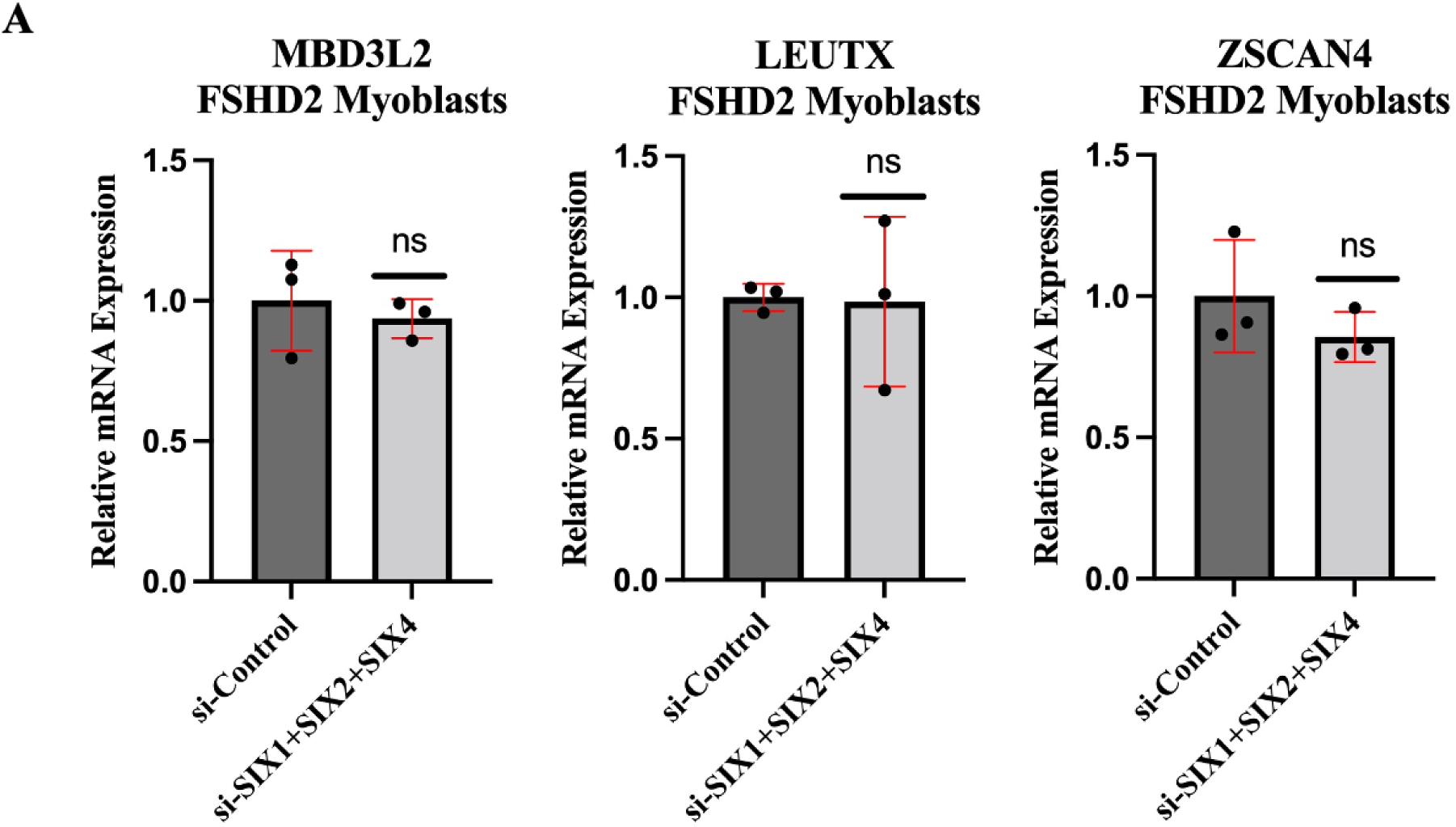
SIX1, SIX2, and SIX4 siRNA knockdown in FSHD2 undifferentiated myoblasts. FSHD2 (MB200) myoblasts were transfected with SIX1, SIX2, SIX4 siRNAs in dual combinations in proliferating myoblasts. (A) qPCR shows the relative mRNA levels DUX4 target genes (*MBD3L2*, *LEUTX* and *ZSCAN4*). Each experimental condition (n=3) was normalized to a negative si-Control with the mean and standard deviation depicted. Asterisks demonstrate statistical significance between the control vs experimental group using an unpaired two tailed *t-*test (ns= not significance; *p<0.05).

**Table S1:** Relative expression levels of each SIX protein in FSHD1 cells. In FSHD1 (54-2) myotubes the Ct Mean and Delta CT (versus internal reference RPL30) were determined from realtime qPCR for each gene of interest: SIX1, SIX2, SIX4 and SIX5 to determine relative expression. The lower the Ct equates to a highly expressed protein. For any Ct >30 the protein mRNA is difficult to detect.

| **Cell Type** | **Gene of Interest** | **CT Mean** | **Delta CT** |
| --- | --- | --- | --- |
| FSHD1 (54-2) | SIX1 | 23.822 | 3.721 |
| FSHD1 (54-2) | SIX2 | 24.186 | 6.178 |
| FSHD1 (54-2) | SIX4 | 24.545 | 4.56 |
| FSHD1 (54-2) | SIX5 | 32.278 | 13.923 |

## References

1. Ahmed, M., J. Xu, and P. X. Xu. 2012. ‘EYA1 and SIX1 drive the neuronal developmental program in cooperation with the SWI/SNF chromatin-remodeling complex and SOX2 in the mammalian inner ear’, Development, 139: 1965–77.

2. Balog, J., P. E. Thijssen, S. Shadle, K. R. Straasheijm, P. J. van der Vliet, Y. D. Krom, M. L. van den Boogaard, A. de Jong, F. Lemmers RJ, R. Tawil, S. J. Tapscott, and S. M. van der Maarel. 2015. ‘Increased DUX4 expression during muscle differentiation correlates with decreased SMCHD1 protein levels at D4Z4’, Epigenetics, 10: 1133–42.

3. Banerji, C. R. S., D. Henderson, R. N. Tawil, and P. S. Zammit. 2020. ‘Skeletal muscle regeneration in facioscapulohumeral muscular dystrophy is correlated with pathological severity’, Hum Mol Genet, 29: 2746–60.

4. Blevins, M. A., C. G. Towers, A. N. Patrick, R. Zhao, and H. L. Ford. 2015. ‘The SIX1-EYA transcriptional complex as a therapeutic target in cancer’, Expert Opin Ther Targets, 19: 213–25.

5. Block, G. J., D. Narayanan, A. M. Amell, L. M. Petek, K. C. Davidson, T. D. Bird, R. Tawil, R. T. Moon, and D. G. Miller. 2013. ‘Wnt/beta-catenin signaling suppresses DUX4 expression and prevents apoptosis of FSHD muscle cells’, Hum Mol Genet, 22: 4661–72.

6. Bosnakovski, D., M. D. Gearhart, E. A. Toso, E. T. Ener, S. H. Choi, and M. Kyba. 2018. ‘Low level DUX4 expression disrupts myogenesis through deregulation of myogenic gene expression’, Sci Rep, 8: 16957.

7. Brouwer, O. F., G. W. Padberg, C. J. Ruys, R. Brand, J. A. de Laat, and J. J. Grote. 1991. ‘Hearing loss in facioscapulohumeral muscular dystrophy’, Neurology, 41: 1878–81.

8. Brouwer, O. F., G. W. Padberg, C. Wijmenga, and R. R. Frants. 1994. ‘Facioscapulohumeral muscular dystrophy in early childhood’, Arch Neurol, 51: 387–94.

9. Campbell, A. E., A. E. Belleville, R. Resnick, S. C. Shadle, and S. J. Tapscott. 2018. ‘Facioscapulohumeral dystrophy: activating an early embryonic transcriptional program in human skeletal muscle’, Hum Mol Genet, 27: R153–R62.

10. Campbell, A. E., J. Oliva, M. P. Yates, J. W. Zhong, S. C. Shadle, L. Snider, N. Singh, S. Tai, Y. Hiramuki, R. Tawil, S. M. van der Maarel, S. J. Tapscott, and F. M. Sverdrup. 2017. ‘BET bromodomain inhibitors and agonists of the beta-2 adrenergic receptor identified in screens for compounds that inhibit DUX4 expression in FSHD muscle cells’, Skelet Muscle, 7: 16.

11. Celegato, B., D. Capitanio, M. Pescatori, C. Romualdi, B. Pacchioni, S. Cagnin, A. Vigano, L. Colantoni, S. Begum, E. Ricci, R. Wait, G. Lanfranchi, and C. Gelfi. 2006. ‘Parallel protein and transcript profiles of FSHD patient muscles correlate to the D4Z4 arrangement and reveal a common impairment of slow to fast fibre differentiation and a general deregulation of MyoD-dependent genes’, Proteomics, 6: 5303–21.

12. Chew, G. L., A. E. Campbell, E. De Neef, N. A. Sutliff, S. C. Shadle, S. J. Tapscott, and R. K. Bradley. 2019. ‘DUX4 Suppresses MHC Class I to Promote Cancer Immune Evasion and Resistance to Checkpoint Blockade’, Dev Cell, 50: 658–71 e7.

13. Cowley, M. V., J. Pruller, M. Ganassi, P. S. Zammit, and C. R. S. Banerji. 2023. ‘An in silico FSHD muscle fiber for modeling DUX4 dynamics and predicting the impact of therapy’, Elife, 12.

14. Cruz, J. M., N. Hupper, L. S. Wilson, J. B. Concannon, Y. Wang, B. Oberhauser, K. Patora-Komisarska, Y. Zhang, D. J. Glass, A. U. Trendelenburg, and B. A. Clarke. 2018. ‘Protein kinase A activation inhibits DUX4 gene expression in myotubes from patients with facioscapulohumeral muscular dystrophy’, J Biol Chem, 293: 11837–49.

15. Davis, T. L., and I. Rebay. 2017. ‘Master regulators in development: Views from the Drosophila retinal determination and mammalian pluripotency gene networks’, Dev Biol, 421: 93–107.

16. De Iaco, A., E. Planet, A. Coluccio, S. Verp, J. Duc, and D. Trono. 2017. ‘DUX-family transcription factors regulate zygotic genome activation in placental mammals’, Nat Genet, 49: 941–45.

17. De Maeyer J, Geese M, Monecke S, Hirsch R, Leng Loke P. 2022. “CASEIN KINASE 1 INHIBITORS FOR USE IN THE TREATMENT OF DISEASES RELATED TO DUX4 EXPRESSION SUCH AS MUSCULAR DYSTROPHY AND CANCER.” In, edited by USPTO. Facio Intellectual Property B.V.

18. Deenen, J. C., H. Arnts, S. M. van der Maarel, G. W. Padberg, J. J. Verschuuren, E. Bakker, S. S. Weinreich, A. L. Verbeek, and B. G. van Engelen. 2014. ‘Population-based incidence and prevalence of facioscapulohumeral dystrophy’, Neurology, 83: 1056–9.

19. Gao, J., D. L. Qin, C. X. Tang, X. Y. Kang, C. J. Song, and C. T. Zhang. 2021. ’Smarcd1 antagonizes the apoptosis of injured MES23.5 DA cells by enhancing the effect of Six2 on GDNF expression’, Neurosci Lett, 760: 136088.

20. Gordon, C. T., S. Xue, G. Yigit, H. Filali, K. Chen, N. Rosin, K. I. Yoshiura, M. Oufadem, T. J. Beck, R. McGowan, A. C. Magee, J. Altmuller, C. Dion, H. Thiele, A. D. Gurzau, P. Nurnberg, D. Meschede, W. Muhlbauer, N. Okamoto, V. Varghese, R. Irving, S. Sigaudy, D. Williams, S. F. Ahmed, C. Bonnard, M. K. Kong, I. Ratbi, N. Fejjal, M. Fikri, S. C. Elalaoui, H. Reigstad, C. Bole-Feysot, P. Nitschke, N. Ragge, N. Levy, G. Tuncbilek, A. S. Teo, M. L. Cunningham, A. Sefiani, H. Kayserili, J. M. Murphy, C. Chatdokmaiprai, A. M. Hillmer, D. Wattanasirichaigoon, S. Lyonnet, F. Magdinier, A. Javed, M. E. Blewitt, J. Amiel, B. Wollnik, and B. Reversade. 2017. ’De novo mutations in SMCHD1 cause Bosma arhinia microphthalmia syndrome and abrogate nasal development’, Nat Genet, 49: 249–55.

21. Goselink, R. J. M., T. H. A. Schreuder, N. van Alfen, I. J. M. de Groot, M. Jansen, Rjlf Lemmers, P. J. van der Vliet, N. van der Stoep, T. Theelen, N. C. Voermans, S. M. van der Maarel, B. G. M. van Engelen, and C. E. Erasmus. 2018. ‘Facioscapulohumeral Dystrophy in Childhood: A Nationwide Natural History Study’, Ann Neurol, 84: 627–37.

22. Gu, Y., Y. Zhao, Y. Zhou, Y. Xie, P. Ju, Y. Long, J. Liu, D. Ni, F. Cao, Z. Lyu, Z. Mao, J. Hao, Y. Li, Q. Wan, Q. Kanyomse, Y. Liu, D. Ren, Y. Ning, X. Li, Q. Zhou, and B. Li. 2016. ‘Zeb1 Is a Potential Regulator of Six2 in the Proliferation, Apoptosis and Migration of Metanephric Mesenchyme Cells’, Int J Mol Sci, 17.

23. Hamanaka, K., D. Sikrova, S. Mitsuhashi, H. Masuda, Y. Sekiguchi, A. Sugiyama, K. Shibuya, Rjlf Lemmers, R. Goossens, M. Ogawa, K. Nagao, C. Obuse, S. Noguchi, Y. K. Hayashi, S. Kuwabara, J. Balog, I. Nishino, and S. M. van der Maarel. 2020. ‘Homozygous nonsense variant in LRIF1 associated with facioscapulohumeral muscular dystrophy’, Neurology, 94: e2441–e47.

24. Hendrickson, P. G., J. A. Dorais, E. J. Grow, J. L. Whiddon, J. W. Lim, C. L. Wike, B. D. Weaver, C. Pflueger, B. R. Emery, A. L. Wilcox, D. A. Nix, C. M. Peterson, S. J. Tapscott, D. T. Carrell, and B. R. Cairns. 2017. ‘Conserved roles of mouse DUX and human DUX4 in activating cleavage-stage genes and MERVL/HERVL retrotransposons’, Nat Genet, 49: 925–34.

25. Himeda, C. L., C. Debarnot, S. Homma, M. L. Beermann, J. B. Miller, P. L. Jones, and T. I. Jones. 2014. ‘Myogenic enhancers regulate expression of the facioscapulohumeral muscular dystrophy-associated DUX4 gene’, Mol Cell Biol, 34: 1942–55.

26. Himeda, C. L., and P. L. Jones. 2019. ‘The Genetics and Epigenetics of Facioscapulohumeral Muscular Dystrophy’, Annu Rev Genomics Hum Genet, 20: 265–91.

27. Himeda, C. L., T. I. Jones, and P. L. Jones. 2015. ‘Facioscapulohumeral muscular dystrophy as a model for epigenetic regulation and disease’, Antioxid Redox Signal, 22: 1463–82.

28. Himeda, C. L., J. A. Ranish, J. C. Angello, P. Maire, R. Aebersold, and S. D. Hauschka. 2004. ‘Quantitative proteomic identification of six4 as the trex-binding factor in the muscle creatine kinase enhancer’, Mol Cell Biol, 24: 2132–43.

29. Hsiao, F. C., A. Williams, E. L. Davies, and I. Rebay. 2001. ‘Eyes absent mediates cross-talk between retinal determination genes and the receptor tyrosine kinase signaling pathway’, Dev Cell, 1: 51–61.

30. Inoue, K., H. Bostan, M. R. Browne, O. F. Bevis, C. D. Bortner, S. A. Moore, A. A. Stence, N. P. Martin, S. H. Chen, A. B. Burkholder, J. L. Li, and N. D. Shaw. 2023. ‘DUX4 double whammy: The transcription factor that causes a rare muscular dystrophy also kills the precursors of the human nose’, Sci Adv, 9: eabq7744.

31. Jagannathan, S. 2022. ‘The evolution of DUX4 gene regulation and its implication for facioscapulohumeral muscular dystrophy’, Biochim Biophys Acta Mol Basis Dis, 1868: 166367.

32. Jagannathan, S., S. C. Shadle, R. Resnick, L. Snider, R. N. Tawil, S. M. van der Maarel, R. K. Bradley, and S. J. Tapscott. 2016. ‘Model systems of DUX4 expression recapitulate the transcriptional profile of FSHD cells’, Hum Mol Genet, 25: 4419–31.

33. Jin, Y., M. Zhang, M. Li, H. Zhang, L. Zhao, C. Qian, S. Li, H. Zhang, M. Gao, B. Pan, R. Li, X. Wan, and C. Cao. 2021. ‘SIX1 Activation Is Involved in Cell Proliferation, Migration, and Anti-inflammation of Acute Ischemia/Reperfusion Injury in Mice’, Front Mol Biosci, 8: 725319.

34. Jones, T. I., J. C. Chen, F. Rahimov, S. Homma, P. Arashiro, M. L. Beermann, O. D. King, J. B. Miller, L. M. Kunkel, C. P. Emerson, Jr., K. R. Wagner, and P. L. Jones. 2012. ‘Facioscapulohumeral muscular dystrophy family studies of DUX4 expression: evidence for disease modifiers and a quantitative model of pathogenesis’, Hum Mol Genet, 21: 4419–30.

35. Jongsma, M. L. M., J. Neefjes, and R. M. Spaapen. 2021. ‘Playing hide and seek: Tumor cells in control of MHC class I antigen presentation’, Mol Immunol, 136: 36–44.

36. Kan, H. E., D. W. Klomp, M. Wohlgemuth, I. van Loosbroek-Wagemans, B. G. van Engelen, G. W. Padberg, and A. Heerschap. 2010. ‘Only fat infiltrated muscles in resting lower leg of FSHD patients show disturbed energy metabolism’, NMR Biomed, 23: 563–8.

37. Karpukhina, A., E. Tiukacheva, C. Dib, and Y. S. Vassetzky. 2021. ‘Control of DUX4 Expression in Facioscapulohumeral Muscular Dystrophy and Cancer’, Trends Mol Med, 27: 588–601.

38. Kumar, J. P. 2009. ‘The sine oculis homeobox (SIX) family of transcription factors as regulators of development and disease’, Cell Mol Life Sci, 66: 565–83.

39. Lassche, S., G. J. Stienen, T. C. Irving, S. M. van der Maarel, N. C. Voermans, G. W. Padberg, H. Granzier, B. G. van Engelen, and C. A. Ottenheijm. 2013. ‘Sarcomeric dysfunction contributes to muscle weakness in facioscapulohumeral muscular dystrophy’, Neurology, 80: 733–7.

40. Le Grand, F., R. Grifone, P. Mourikis, C. Houbron, C. Gigaud, J. Pujol, M. Maillet, G. Pages, M. Rudnicki, S. Tajbakhsh, and P. Maire. 2012. ‘Six1 regulates stem cell repair potential and self-renewal during skeletal muscle regeneration’, J Cell Biol, 198: 815–32.

41. Lemmers, R. J., P. J. van der Vliet, R. Klooster, S. Sacconi, P. Camano, J. G. Dauwerse, L. Snider, K. R. Straasheijm, G. J. van Ommen, G. W. Padberg, D. G. Miller, S. J. Tapscott, R. Tawil, R. R. Frants, and S. M. van der Maarel. 2010. ‘A unifying genetic model for facioscapulohumeral muscular dystrophy’, Science, 329: 1650–3.

42. Li, L., Y. Liang, L. Kang, Y. Liu, S. Gao, S. Chen, Y. Li, W. You, Q. Dong, T. Hong, Z. Yan, S. Jin, T. Wang, W. Zhao, H. Mai, J. Huang, X. Han, Q. Ji, Q. Song, C. Yang, S. Zhao, X. Xu, and Q. Ye. 2018. ‘Transcriptional Regulation of the Warburg Effect in Cancer by SIX1’, Cancer Cell, 33: 368–85 e7.

43. Lutz, K. L., L. Holte, S. A. Kliethermes, C. Stephan, and K. D. Mathews. 2013. ‘Clinical and genetic features of hearing loss in facioscapulohumeral muscular dystrophy’, Neurology, 81: 1374–7.

44. Maire, P., M. Dos Santos, R. Madani, I. Sakakibara, C. Viaut, and M. Wurmser. 2020. ‘Myogenesis control by SIX transcriptional complexes’, Semin Cell Dev Biol, 104: 51–64.

45. Meurer, L., L. Ferdman, B. Belcher, and T. Camarata. 2021. ‘The SIX Family of Transcription Factors: Common Themes Integrating Developmental and Cancer Biology’, Front Cell Dev Biol, 9: 707854.

46. Mocciaro, E., V. Runfola, P. Ghezzi, M. Pannese, and D. Gabellini. 2021. ‘DUX4 Role in Normal Physiology and in FSHD Muscular Dystrophy’, Cells, 10.

47. Moody, S. A., and A. S. LaMantia. 2015. ‘Transcriptional regulation of cranial sensory placode development’, Curr Top Dev Biol, 111: 301–50.

48. Oliva, J., S. Galasinski, A. Richey, A. E. Campbell, M. J. Meyers, N. Modi, J. W. Zhong, R. Tawil, S. J. Tapscott, and F. M. Sverdrup. 2019. ‘Clinically Advanced p38 Inhibitors Suppress DUX4 Expression in Cellular and Animal Models of Facioscapulohumeral Muscular Dystrophy’, J Pharmacol Exp Ther, 370: 219–30.

49. Padberg, G. W., O. F. Brouwer, R. J. de Keizer, G. Dijkman, C. Wijmenga, J. J. Grote, and R. R. Frants. 1995. ‘On the significance of retinal vascular disease and hearing loss in facioscapulohumeral muscular dystrophy’, Muscle Nerve Suppl: S73–80.

50. Park, B. Y., and J. P. Saint-Jeannet. 2010. Induction and Segregation of the Vertebrate Cranial Placodes (San Rafael (CA)).

51. Pineda, J. M. B., and R. K. Bradley. 2023. ‘DUX4 is a common driver of immune evasion and immunotherapy failure in metastatic cancers’, bioRxiv.

52. Rickard, A. M., L. M. Petek, and D. G. Miller. 2015. ‘Endogenous DUX4 expression in FSHD myotubes is sufficient to cause cell death and disrupts RNA splicing and cell migration pathways’, Hum Mol Genet, 24: 5901–14.

53. Rickard, A.; Schmidt, U.; Kiselyov, A. 2023. “Inhibitors of DUX4 induction for regulation of muscle function.” In. US: Sonic Master Limited.

54. Rojas, L. A., E. Valentine, A. Accorsi, J. Maglio, N. Shen, A. Robertson, S. Kazmirski, P. Rahl, R. Tawil, D. Cadavid, L. A. Thompson, L. Ronco, A. N. Chang, A. M. Cacace, and O. Wallace. 2020. ‘p38alpha Regulates Expression of DUX4 in a Model of Facioscapulohumeral Muscular Dystrophy’, J Pharmacol Exp Ther, 374: 489–98.

55. Ruf, R. G., P. X. Xu, D. Silvius, E. A. Otto, F. Beekmann, U. T. Muerb, S. Kumar, T. J. Neuhaus, M. J. Kemper, R. M. Raymond, Jr., P. D. Brophy, J. Berkman, M. Gattas, V. Hyland, E. M. Ruf, C. Schwartz, E. H. Chang, R. J. Smith, C. A. Stratakis, D. Weil, C. Petit, and F. Hildebrandt. 2004. ‘SIX1 mutations cause branchio-oto-renal syndrome by disruption of EYA1-SIX1-DNA complexes’, Proc Natl Acad Sci U S A, 101: 8090–5.

56. Sakakibara, I., M. Wurmser, M. Dos Santos, M. Santolini, S. Ducommun, R. Davaze, A. Guernec, K. Sakamoto, and P. Maire. 2016. ‘Six1 homeoprotein drives myofiber type IIA specialization in soleus muscle’, Skelet Muscle, 6: 30.

57. Segales, J., A. B. Islam, R. Kumar, Q. C. Liu, P. Sousa-Victor, F. J. Dilworth, E. Ballestar, E. Perdiguero, and P. Munoz-Canoves. 2016. ‘Chromatin-wide and transcriptome profiling integration uncovers p38alpha MAPK as a global regulator of skeletal muscle differentiation’, Skelet Muscle, 6: 9.

58. Seo, H. C., J. Curtiss, M. Mlodzik, and A. Fjose. 1999. ‘Six class homeobox genes in drosophila belong to three distinct families and are involved in head development’, Mech Dev, 83: 127–39.

59. Shaw, N. D., H. Brand, Z. A. Kupchinsky, H. Bengani, L. Plummer, T. I. Jones, S. Erdin, K. A. Williamson, J. Rainger, A. Stortchevoi, K. Samocha, B. B. Currall, D. S. Dunican, R. L. Collins, J. R. Willer, A. Lek, M. Lek, M. Nassan, S. Pereira, T. Kammin, D. Lucente, A. Silva, C. M. Seabra, C. Chiang, Y. An, M. Ansari, J. K. Rainger, S. Joss, J. C. Smith, M. F. Lippincott, S. S. Singh, N. Patel, J. W. Jing, J. R. Law, N. Ferraro, A. Verloes, A. Rauch, K. Steindl, M. Zweier, I. Scheer, D. Sato, N. Okamoto, C. Jacobsen, J. Tryggestad, S. Chernausek, L. A. Schimmenti, B. Brasseur, C. Cesaretti, J. E. Garcia-Ortiz, T. P. Buitrago, O. P. Silva, J. D. Hoffman, W. Muhlbauer, K. W. Ruprecht, B. L. Loeys, M. Shino, A. M. Kaindl, C. H. Cho, C. C. Morton, R. R. Meehan, V. van Heyningen, E. C. Liao, R. Balasubramanian, J. E. Hall, S. B. Seminara, D. Macarthur, S. A. Moore, K. I. Yoshiura, J. F. Gusella, J. A. Marsh, J. M. Graham, Jr., A. E. Lin, N. Katsanis, P. L. Jones, W. F. Crowley, Jr., E. E. Davis, D. R. FitzPatrick, and M. E. Talkowski. 2017. ‘SMCHD1 mutations associated with a rare muscular dystrophy can also cause isolated arhinia and Bosma arhinia microphthalmia syndrome’, Nat Genet, 49: 238–48.

60. Sidlauskaite, E., L. Le Gall, V. Mariot, and J. Dumonceaux. 2020. ‘DUX4 Expression in FSHD Muscles: Focus on Its mRNA Regulation’, J Pers Med, 10.

61. Simone, C., S. V. Forcales, D. A. Hill, A. N. Imbalzano, L. Latella, and P. L. Puri. 2004. ‘p38 pathway targets SWI-SNF chromatin-remodeling complex to muscle-specific loci’, Nat Genet, 36: 738–43.

62. Smith, A. A., Y. Nip, S. R. Bennett, D. C. Hamm, Rjlf Lemmers, P. J. van der Vliet, M. Setty, S. M. van der Maarel, and S. J. Tapscott. 2023. ‘DUX4 expression in cancer induces a metastable early embryonic totipotent program’, Cell Rep, 42: 113114.

63. Snider, L., L. N. Geng, R. J. Lemmers, M. Kyba, C. B. Ware, A. M. Nelson, R. Tawil, G. N. Filippova, S. M. van der Maarel, S. J. Tapscott, and D. G. Miller. 2010. ‘Facioscapulohumeral dystrophy: incomplete suppression of a retrotransposed gene’, PLoS Genet, 6: e1001181.

64. Soni, U. K., K. Roychoudhury, and R. S. Hegde. 2021. ‘The Eyes Absent proteins in development and in developmental disorders’, Biochem Soc Trans, 49: 1397–408.

65. Statland, J., and R. Tawil. 2014. ‘Facioscapulohumeral muscular dystrophy’, Neurol Clin, 32: 721–8, ix.

66. Talbot, J., and L. Maves. 2016. ‘Skeletal muscle fiber type: using insights from muscle developmental biology to dissect targets for susceptibility and resistance to muscle disease’, Wiley Interdiscip Rev Dev Biol, 5: 518–34.

67. Tassin, A., D. Laoudj-Chenivesse, C. Vanderplanck, M. Barro, S. Charron, E. Ansseau, Y. W. Chen, J. Mercier, F. Coppee, and A. Belayew. 2013. ‘DUX4 expression in FSHD muscle cells: how could such a rare protein cause a myopathy?’, J Cell Mol Med, 17: 76–89.

68. Tawil, R., and S. M. Van Der Maarel. 2006. ‘Facioscapulohumeral muscular dystrophy’, Muscle Nerve, 34: 1–15.

69. Tawil, R., S. M. van der Maarel, and S. J. Tapscott. 2014. ‘Facioscapulohumeral dystrophy: the path to consensus on pathophysiology’, Skelet Muscle, 4: 12.

70. Tihaya, M. S., K. Mul, J. Balog, J. C. de Greef, S. J. Tapscott, R. Tawil, J. M. Statland, and S. M. van der Maarel. 2023. ‘Facioscapulohumeral muscular dystrophy: the road to targeted therapies’, Nat Rev Neurol, 19: 91–108.

71. van der Maarel, S. M., R. R. Frants, and G. W. Padberg. 2007. ‘Facioscapulohumeral muscular dystrophy’, Biochim Biophys Acta, 1772: 186–94.

72. van der Maarel, S. M., D. G. Miller, R. Tawil, G. N. Filippova, and S. J. Tapscott. 2012. ‘Facioscapulohumeral muscular dystrophy: consequences of chromatin relaxation’, Curr Opin Neurol, 25: 614–20.

73. van der Maarel, S. M., R. Tawil, and S. J. Tapscott. 2011. ‘Facioscapulohumeral muscular dystrophy and DUX4: breaking the silence’, Trends Mol Med, 17: 252–8.

74. van Overveld, P. G., R. J. Lemmers, L. A. Sandkuijl, L. Enthoven, S. T. Winokur, F. Bakels, G. W. Padberg, G. J. van Ommen, R. R. Frants, and S. M. van der Maarel. 2003. ‘Hypomethylation of D4Z4 in 4q-linked and non-4q-linked facioscapulohumeral muscular dystrophy’, Nat Genet, 35: 315–7.

75. Viaut, C., S. Weldon, and A. Munsterberg. 2021. ‘Fine-tuning of the PAX-SIX-EYA-DACH network by multiple microRNAs controls embryo myogenesis’, Dev Biol, 469: 68–79.

76. Wijmenga, C., R. R. Frants, O. F. Brouwer, P. Moerer, J. L. Weber, and G. W. Padberg. 1990. ‘Location of facioscapulohumeral muscular dystrophy gene on chromosome 4’, Lancet, 336: 651–3.

77. Wu, W., Z. Ren, P. Li, D. Yu, J. Chen, R. Huang, and H. Liu. 2015. ‘Six1: a critical transcription factor in tumorigenesis’, Int J Cancer, 136: 1245–53.

78. Wurmser, M., R. Madani, N. Chaverot, S. Backer, M. Borok, M. Dos Santos, G. Comai, S. Tajbakhsh, F. Relaix, M. Santolini, R. Sambasivan, R. Jiang, and P. Maire. 2023. ‘Overlapping functions of SIX homeoproteins during embryonic myogenesis’, PLoS Genet, 19: e1010781.

79. Yajima, H., N. Motohashi, Y. Ono, S. Sato, K. Ikeda, S. Masuda, E. Yada, H. Kanesaki, Y. Miyagoe-Suzuki, S. Takeda, and K. Kawakami. 2010. ‘Six family genes control the proliferation and differentiation of muscle satellite cells’, Exp Cell Res, 316: 2932–44.

80. Yao, Z., L. Snider, J. Balog, R. J. Lemmers, S. M. Van Der Maarel, R. Tawil, and S. J. Tapscott. 2014. ‘DUX4- induced gene expression is the major molecular signature in FSHD skeletal muscle’, Hum Mol Genet, 23: 5342–52.

81. Zetser, A., E. Gredinger, and E. Bengal. 1999. ‘p38 mitogen-activated protein kinase pathway promotes skeletal muscle differentiation. Participation of the Mef2c transcription factor’, J Biol Chem, 274: 5193–200.

